# Kidney disease reprograms microbiome-host signaling to promote heart failure

**DOI:** 10.64898/2026.01.10.698142

**Authors:** Alex Yarritu, Wibke Anders, Arne Thiele, Olena Potapenko, Fabian Schumacher, István A. Szijartó, Ariana Matz-Rauch, Jakob Versnjak, Natnael Gebremedhin, Victoria McParland, Simon Heckscher, Sakshi Kamboj, Giuseppe Trimarchi, Harithaa Anandakumar, Franziska Fuckert, Carina Hoffmann, Sara A. Hassan, Paul M. Bonnekoh, Moritz I. Wimmer, Felix Behrens, Jakob Voelkl, Rafael Kramann, Ulrich Kintscher, Titus Kuehne, Burkhard Kleuser, Alma Zernecke, Peter J. Oefner, Wolfram Gronwald, Katja Dettmer, Kai-Uwe Eckardt, Dominik N. Müller, Marcus Kelm, Johannes Holle, Hendrik Bartolomaeus, Nicola Wilck

**Author notes:** Correspondence: Nicola Wilck, Experimental and Clinical Research Center, Lindenberger Weg 80, 13125 Berlin. These authors contributed equally to this work (Alex Yarritu, Wibke Anders). These authors contributed equally to this work (Hendrik Bartolomaeus, Nicola Wilck).

## Abstract

**Background:** Heart failure is prevalent in chronic kidney disease (CKD) and linked to chronic inflammation. CKD-typical gut microbiome dysbiosis may stimulate inflammation, as bacterial aromatic metabolites are highly abundant and engage transcriptional programs through the aryl hydrocarbon receptor (AhR). Whether this axis drives cardiac remodeling and is therapeutically targetable remains unknown.

**Methods:** We used the subtotal nephrectomy model (STNx) and microbiome depletion by oral antibiotics. We investigated cardiac and renal function, AhR activity, metabolite profiles, and immunophenotypes by flow cytometry and transcriptomics. Candidate metabolite indoxyl sulfate (IxS) was tested in experimental HFpEF. In vivo and in vitro AhR inhibition (AhRi) was performed using a clinically tested compound. Mechanistic studies were performed in primary human and murine cardiac fibroblasts and T cells, as well as translational validation using UK Biobank data.

**Results:** Microbiome depletion lowered bacterial metabolites and attenuated cardiac fibrosis and diastolic dysfunction in STNx, identifying AhR-driven expansion of interleukin-17A (IL-17A)-producing T helper cells (T_H_17) as key effector. Plasma IL-17A was stage-dependently elevated in CKD patients, particularly in HFpEF, and associated with all-cause mortality. Bacterial metabolite IxS promoted T_H_17 polarization and exacerbated cardiac dysfunction in HFpEF. AhRi using a small molecule inhibitor reduced T_H_17 abundance and attenuated cardiac fibrosis in STNx. Mechanistically, AhR and IL-17A signaling synergistically induced a conserved pro-fibrotic phenotype in human and murine cardiac fibroblasts, and AhR inhibition blocked ECM production in response to CKD patient serum.

**Conclusion:** A microbiome-AhR-IL-17A axis drives CKD-associated cardiac fibrosis. AhRi prevents remodeling, highlighting a potential therapeutic avenue to prevent cardiorenal multimorbidity.

## Introduction

Cardiovascular disease (CVD) is the most common comorbidity and the primary cause of mortality in chronic kidney disease (CKD)^1^. CKD is a global health burden of increasing relevance^2^, and adverse cardiac remodeling very common in these patients. Heart failure (HF) is particularly prevalent in CKD, especially with preserved ejection fraction (HFpEF)^3^. The high cardiovascular burden in CKD is driven by a persistent low-grade inflammatory state^4^, which has been linked to an altered gut microbiome composition and function by us^5^ and others^6^. Importantly, CKD impairs the urinary clearance of microbiome-derived metabolites, leading to their accumulation and partly toxic effects^7,8^. Immune cells are increasingly recognized as active mediators of cardiac injury^9^. T cells play a causal role in mediating diastolic dysfunction in experimental CKD^10^, implicating adaptive immunity as a causal contributor. However, the CKD-specific cues licensing T cell pathogenicity as well as the effector mechanisms through which they promote cardiac remodeling remain incompletely defined.

Although the gut microbiome strongly influences adaptive immunity, it is not yet targeted by standard CKD therapies. CKD is characterized by altered production of microbiome-derived metabolites, including increased generation of ligands for the aryl hydrocarbon receptor (AhR), a widely expressed host transcription factor^11^. Indoxyl sulfate (IxS), a prototypical uremic toxin and agonistic ligand of the AhR, is generated from gut microbial metabolism of dietary tryptophan followed by hepatic sulfation. Upon ligand binding and activation, the AhR translocates to the nucleus, where it regulates gene expression and immune cell differentiation^12^. In CD4^+^ T helper cells (T_H_), AhR signaling influences T_H_ cell fate, biasing differentiation toward T_H_17, regulatory T cells (T_reg_), or other lineages depending on ligand type, concentration, and cellular context^13^, thereby shaping the inflammatory microenvironment^14^. Interleukin-17A (IL-17A), the principal effector cytokine of T_H_17 cells, is a mediator of autoimmune and inflammatory diseases, where neutralizing antibodies are in clinical use. While IL-17A has been implicated in cardiac inflammation in kidney-independent scenarios such as autoimmune myocarditis^15^, its role in CKD-associated cardiac remodeling has not been defined.

Here, we examine how gut microbiome-immune interactions drive adverse cardiac remodeling in kidney disease. We show that microbiome depletion confers cardiac protection and identify IL-17A as a microbiome-dependent driver of cardiac fibrosis and dysfunction in CKD. IL-17A is increased in CKD patients and even more so with coexisting HFpEF. Bacterial metabolite IxS promotes T_H_17 differentiation via the AhR *in vitro* and induces T_H_17 and aggravates cardiac remodeling in experimental HFpEF. Importantly, AhR inhibition suppresses T_H_17 induction and cardiac fibrosis despite metabolite accumulation. We identify murine and human cardiac fibroblasts as AhR-modulated, IL-17A-responsive effectors. These findings reveal microbiome-dependent, pro-fibrotic pathways and position the AhR as a therapeutic target for cardiovascular protection in CKD.

## Methods

Extended methods are provided in the supplementary material.

### Data and code availability

All code, data, and scripts used in this study will be made publicly available. The code and scripts will be made accessible on Zenodo. The raw sequencing data will be deposited in the NCBI Sequence Read Archive.

### Ethics

All animal experiments were performed in accordance with the German/European law for animal protection. Experimental protocols were approved by the local ethic authorities (LaGeSo) under the G0109/21 and G0019/21 licenses.

Patients with CKD stage 5 dependent on hemodialysis were recruited Department of Nephrology and Medical Intensive Care at Charité – Universitätsmedizin Berlin. Healthy volunteers were recruited at the Experimental and Clinical Research Center (ECRC) of Charité – Universitätsmedizin Berlin. Ethical approval was obtained from the ethics committee of Charité – Universitätsmedizin Berlin (EA2/162/17) and all participants provided written informed consent before biomaterial was collected.

Ethical approval for UK Biobank was granted by the North West Multi-Centre Research Ethics Committee (reference 11/NW/0382), and all participants provided written informed consent.

### Statistical analysis

Data are presented as Mean ± SEM with individual datapoints. Normality distribution of data was assessed using Q-Q-plots before statistical comparisons. For four-group comparisons, statistical significance was assessed by ANOVA or Kruskal-Wallis test, followed by Tukey’s or Dunn’s *post hoc* correction, as appropriate. For two-group comparisons, statistical significance was assessed by Student’s t-test or Mann-Whitney U test.

Metabolome analysis was performed using Mann-Whitney U test and adjusted using the false discovery rate (Benjamini-Hochberg, FDR). Microbiome analysis was performed using Mann-Whitney U test for alpha diversity indices and differential abundances were assessed using the Brunner-Munzel test with FDR correction. Bulk RNA sequencing was analyzed using a generalized linear model with FDR correction implemented in DeSeq2. Gene set enrichment analyses were performed using Fisher’s exact test followed by FDR correction. Single-cell RNA sequencing datasets were analyzed using Seurat. Differential gene expression was performed within the MAST framework, and cytokine response enrichment was assessed using the Immune Response Enrichment Analysis (IREA) software.

UK Biobank analysis was performed in Python 3.11. Survival analysis was performed using log-rank test, and Cox proportional hazards model and hazard ratios with 95% confidence intervals were reported. Normality of continuous variables was assessed using the Shapiro-Wilk test. Group differences were evaluated using Welch’s t-test or Mann-Whitney U test, as appropriate. For comparison of multiple groups, FDR procedure was used.

## Results

### Gut microbiota accelerates cardiac remodeling in kidney disease

To assess the contribution of the gut microbiota to the host’s response to kidney disease, we compared colonized and microbiome-depleted (oral antibiotics, Abx) 129/Sv mice subjected to subtotal nephrectomy (STNx), with sham-operated mice (Sham) as controls (Fig.1a). Disease progression was monitored over a 13-week period. Reduced 16S rRNA gene copies in cecal content confirmed successful microbiota depletion over the course of the experiment (Fig.S1a) and cecal weight was increased in Abx-treated mice, reminiscent of the germ-free (GF) mouse cecum (Fig.S1b)^16^. 16S rRNA gene sequencing of cecal samples from the non-microbiome-depleted groups showed a minor reduction in alpha diversity (Fig.S1c-e) in STNx mice while phyla composition remained similar (Fig.S1f). Principal component analysis (PCA) indicated a separation of bacterial communities (Fig.S1g). We found this to be due to an increase in known CKD-associated genera such as *Bacteroides* and *Ruminococcus* (Fig.S1h)^17^. Notably, the taxonomic shifts observed after STNx were less pronounced than those reported in patients with CKD^18^, likely because factors such as diet and medication - major drivers of microbial variation in humans - are standardized or absent in our mouse model^19^. Functionally, bacterial metabolites are particularly relevant in kidney disease, as they can accumulate as a function of increased microbial production and insufficient renal excretion^8^. Thus, we performed plasma metabolomics by nuclear magnetic resonance (NMR), covering a large number of metabolites (Fig.1b). Antibiotic depletion led to a substantial reduction of metabolite accumulation in STNx mice, with most pronounced changes in the aromatic region of the NMR spectrum (Fig.1c). We therefore quantified a targeted panel of aromatic metabolites in plasma by mass spectrometry (MS, Fig.1d). Bacterially derived toxins such as indoxyl sulfate (IxS), p-cresol sulfate and phenyl sulfate were significantly increased in STNx (Fig.1e) and consequently reduced by Abx treatment. Thus, kidney disease induces taxonomic alterations in the microbiome and elevates bacterial metabolite concentrations in plasma, with both effects abrogated by microbiome depletion.

**Figure 1.**
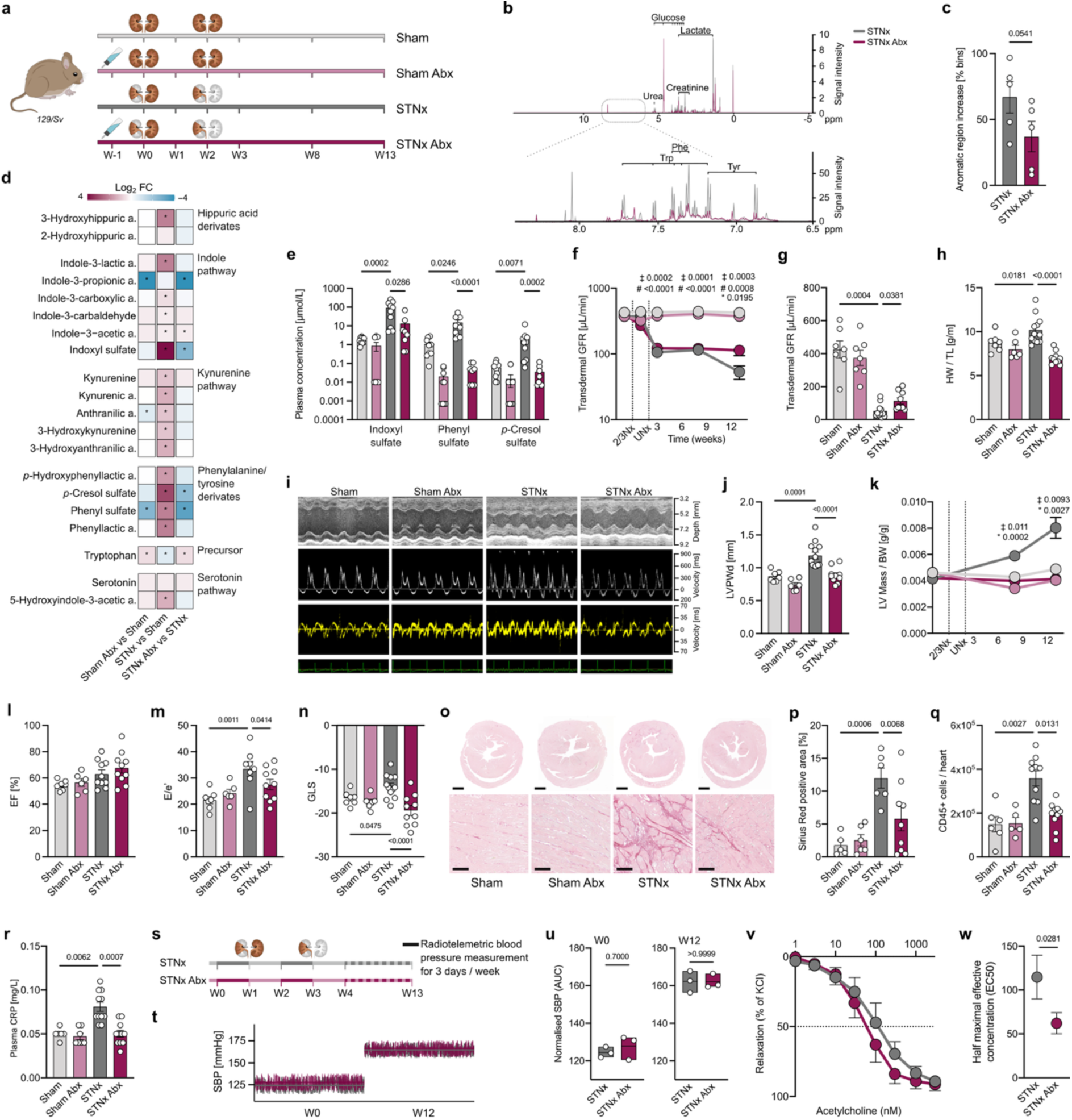
Depletion of microbiome-derived metabolites attenuates cardiac remodeling in mice with impaired kidney function. **(a)** 129/Sv mice were randomized to receive a broad-spectrum, oral antibiotic cocktail (Abx) or control drinking water. After one week, mice underwent two-stage subtotal nephrectomy (STNx) or sham surgery. **(b)** Representative ^1^H-NMR plasma spectra 13 weeks after surgery and **(c)** determination of summed-up peak areas within the aromatic region, normalized to the respective sham group. **(d)** Heatmap of plasma metabolites quantified by liquid chromatography-mass spectrometry (LC-MS). **(e)** Plasma concentrations of sulfated microbiome-derived metabolites quantified by LC-MS. Transdermal glomerular filtration rate (GFR) normalized measured **(f)** longitudinally and **(g)** at endpoint. **(h)** Heart weight normalized to tibia length at the endpoint. **(i)** Representative echocardiographic images at the endpoint. **(j)** Left ventricular posterior wall thickness at end-diastole (LVPWd). **(k)** Longitudinal changes in normalized left ventricular (LV) mass. **(l)** Ejection fraction (EF), **(m)** E/e′ ratio, **(n)** global longitudinal strain (GLS) at the endpoint. **(o)** Histological assessment of left ventricular fibrosis by Sirius Red staining (scale bars: upper, 1 mm; lower, 100 µm) and **(p)** quantification of fibrotic area. **(q)** Quantification of leukocytes in the heart at the endpoint by flow cytometry. **(r)** Plasma C-reactive protein (CRP) levels at the endpoint. **(s)** Radiotelemetric blood pressure (BP) measurement. **(t)** Representative average systolic BP (SBP) plots at baseline (week 0) and week 12, and **(u)** quantification. **(v)** Acetylcholine (ACh)-induced endothelial-dependent vasorelaxation of mesenteric arteries assessed by wire myography, and **(w)** corresponding EC₅₀ values. Symbols in **(f)** and **(k)** indicate the following comparisons: Sham vs. STNx (‡), Sham Abx vs. STNx Abx (#), and STNx vs. STNx Abx (*). Data are shown as mean ± SEM. Each dot represents one mouse. Statistical analyses were performed using one-way ANOVA or Kruskal-Wallis test followed by Tukey’s or Dunn’s *post hoc* correction, as appropriate, unless otherwise indicated. Mann Whitney U-tests with false discovery rate correction according to Benjamini-Hochberg were used for metabolomics data. Two-group comparisons for the aromatic NMR region **(c)** and EC₅₀ values **(w)** were performed using unpaired two-tailed Student’s t-test. Two-group comparisons of systolic blood pressure in **(u)** were assessed by Mann-Whitney U test.

To assess the impact of microbiome depletion on the degree of kidney dysfunction, we measured glomerular filtration rate (GFR) longitudinally. While both STNx groups displayed a comparable initial GFR reduction until week 8 (Fig.1f), Abx-treated STNx mice preserved higher GFR at the endpoint (Fig.1g) despite similar remnant kidney mass (Fig.S2a). Measurement of plasma cystatin C (CysC), an endogenous filtration marker, confirmed this observation (Fig.S2b).

Focusing on cardiovascular comorbidities as the leading cause of mortality in CKD^1,20^, we next assessed cardiac and vascular damage. STNx induced cardiac remodeling as reflected by increased heart weight, which was abolished by microbiome depletion (Fig.1h). Longitudinal echocardiographic assessment of cardiac dimensions corroborated this effect, becoming evident from week 8 onward (Fig.1i-k; Table S1) despite similar GFR at this time point. While systolic function (ejection fraction, EF) was preserved in all groups (Fig. 1l; Table S1), STNx mice displayed an increase in E/e’ (Fig.1i, m; Table S1) indicative of diastolic dysfunction, and a reduction in global longitudinal strain (GLS, Fig.1n; Table S1), reflecting ventricular stiffening - both rescued by microbiome depletion. Adverse cardiac remodeling is characterized by cardiomyocyte hypertrophy, fibrosis and inflammation^21^. Interestingly, we observed comparable cardiomyocyte sizes in STNx (Fig.S2c-d), suggesting that the increased heart weight is driven by extracellular matrix (ECM) deposition. Sirius Red staining revealed increased collagen deposition in STNx hearts, which was mitigated in microbiome-depleted STNx mice (Fig.1o-p). In addition, STNx mice exhibited increased cardiac leukocyte infiltration (Fig.1q), comprising mostly neutrophils and T cells (Fig.S2e), which was ameliorated in Abx-treated STNx mice. In agreement, plasma C-reactive protein (CRP) - a routine clinical marker of systemic inflammation associated with cardiovascular risk^22^ - was elevated in STNx mice and reduced by Abx treatment (Fig.1r). These findings demonstrate that microbiome depletion prevents hallmarks of systemic inflammation, adverse cardiac remodeling and diastolic dysfunction induced by kidney disease.

Reduced kidney function typically leads to hypertension, a common cause for adverse cardiac remodeling. Further evidence suggests that the gut microbiota influences blood pressure (BP) and cardiac remodeling^23,24^. We therefore measured BP longitudinally by radiotelemetry in both control and Abx-treated STNx mice (Fig.1s). Both groups exhibited a similar increase in both systolic BP (SBP) and diastolic BP (DBP) throughout the experiment, which was sustained towards the study endpoint (Fig.1t-u, S2f-i), suggesting that the cardioprotective effects observed in microbiota-depleted STNx mice are independent of BP. To assess vascular function, we measured vasorelaxation in isolated mesenteric arteries. Arteries from Abx-treated STNx mice displayed an improved endothelium-dependent relaxation compared to those from STNx mice (Fig.1v-w). Taken together, microbiome depletion protected mice with reduced kidney function from further GFR loss, inflammation, cardiac and vascular dysfunction despite a similar blood pressure increase, suggesting these benefits are linked to immunological mechanisms.

### Microbiome-Dependent T_H_17 Immunity and AhR Activation

Since cardiac leukocyte influx was attenuated by microbiome depletion and T cells have been shown to causally contribute to CKD-associated cardiac dysfunction^10^, we hypothesized that microbiome-immune interactions, and microbiome-T cell interaction in particular, shape the pathological cardiac response to STNx. We performed flow cytometry on splenic leukocytes with a specific focus on T cells (Fig.2a). STNx induced a T cell shift towards CD4^+^ T_H_ subsets (Fig.2b), which was prevented by Abx treatment. Within the T_H_ cell compartment, CD44^+^CD62L^−^ effector memory cells expanded in both FoxP3^−^ conventional T_H_ cells (T_conv_, Fig.2c) and FoxP3^+^ T_reg_ (Fig.2d), again abrogated by Abx. Moreover, splenic T_H_ cells from STNx mice showed an increased abundance of RORγt^+^ T_conv_ (T_H_17, Fig.2e) and RORγt^+^ T_reg_, hereafter referred to as T_reg_17 cells (Fig.2f), both prevented by Abx. Similar patterns of T_H_17 expansion were detected in circulating and intestinal lamina propria T cells (Fig.2g), indicating a microbiome-dependent T_H_17-type inflammatory response in experimental CKD. Consistent with the known T_H_17-neutrophil interaction^25^, T_H_17 patterns were mirrored in neutrophil abundance (Fig.S3a and Fig.2h).

**Figure 2.**
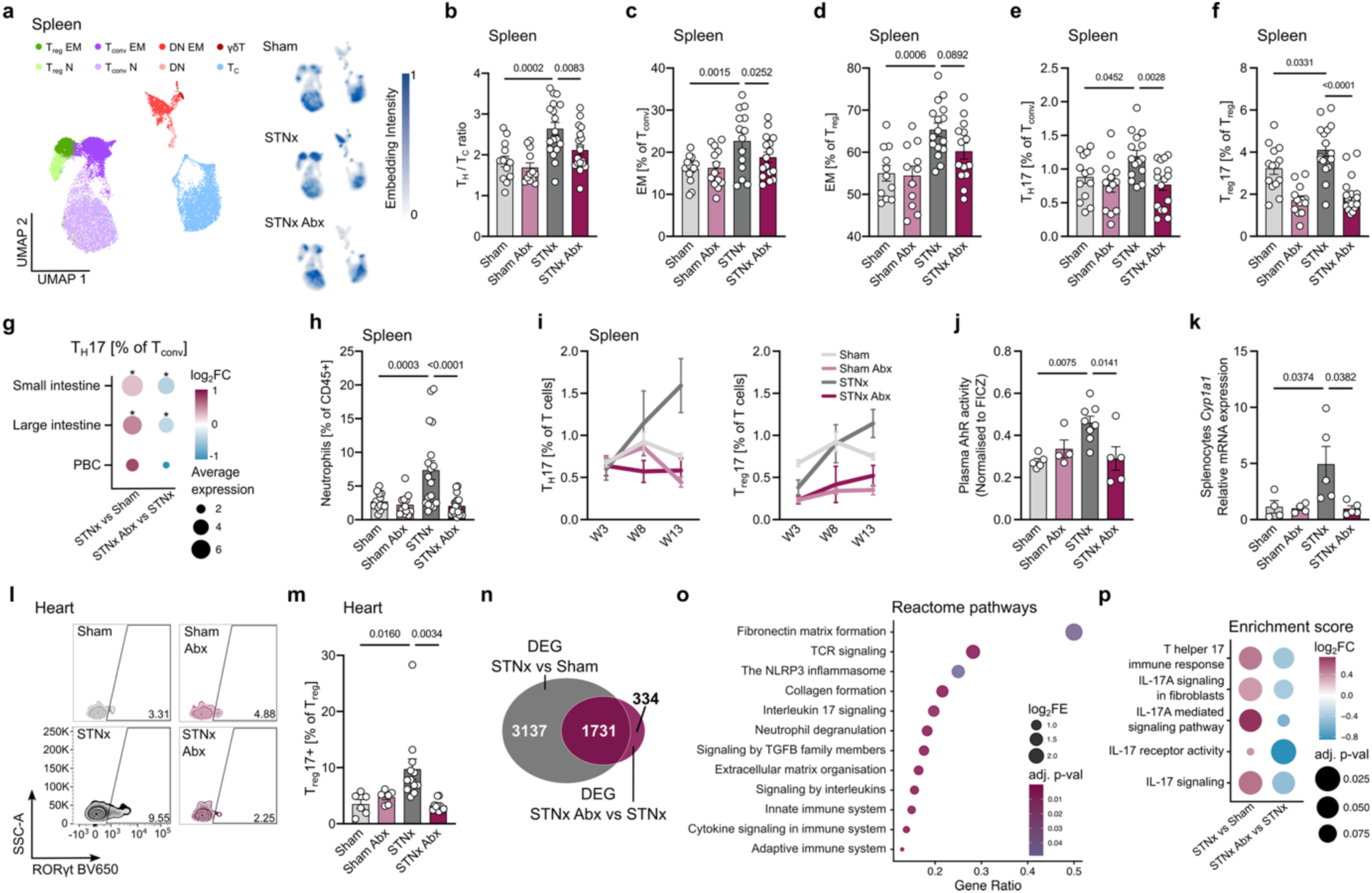
Kidney disease induces microbiota-dependent T_H_17-driven systemic and cardiac inflammation. Immune cells were isolated from spleens and analyzed by flow cytometry. **(a)** UMAP projection of splenic T cell populations based on flow cytometry. **(b)** Ratio of T helper (T_H_, CD45^+^ CD3^+^ TCRγδ^−^ CD4⁺ CD8^−^) to cytotoxic T cells (T_C_, CD45^+^ CD3^+^ TCRγδ^−^ CD4⁺ CD8^−^). **(c)** Frequencies of effector memory (EM, CD44^+^ CD62L^−^) conventional T cells (T_conv_, FoxP3^−^ T_H_). **(d)** Frequencies of EM regulatory T cells (T_reg_, FoxP3^+^ T_H_). **(e)** Frequencies of RORγt⁺ T_conv_. **(f)** Frequencies of RORγt⁺ T_reg_. **(g)** Changes in RORγt⁺ T_conv_ abundance between experimental groups (* p-adj. < 0.1). **(h)** Abundance of neutrophils (CD45^+^ CD3^−^ NK1.1^−^ CD11b^+^ Ly6G^+^) among CD45⁺ cells. **(i)** Longitudinal frequencies of RORγt⁺ T_conv_ (left) and T_reg_ (right). **(j)** Aryl hydrocarbon receptor (AhR) activation of plasma in reporter cell line. **(k)** *Cyp1a1* expression in splenocytes. **(l)** Representative flow cytometric plots of cardiac RORγt⁺ T_reg_ (T_reg_17) and **(m)** quantification. **(n)** Overlap of differentially expressed genes (DEG) in cardiac bulk mRNA-sequencing between the two indicated comparisons. **(o)** Significantly enriched biological pathways in the cardiac transcriptome. **(p)** Targeted gene ontology (GO) and pathway analysis of IL-17A-related differentially enriched gene sets. Data are shown as mean ± SEM. Each dot represents one mouse. Statistical analyses were performed using one-way ANOVA or Kruskal-Wallis test followed by Tukey’s or Dunn’s *post hoc* correction, as appropriate, unless otherwise indicated. For **(g)**, Benjamini-Hochberg false discovery rate (FDR) correction was performed. For RNA sequencing, DEG were generated using the DeSeq2 pipeline with appropriate FDR correction. Gene set enrichment analyses were performed using Fisher’s exact test followed by FDR correction.

While T_reg_17 cells conventionally restrain mucosal inflammation^26^, they can also adopt T_H_17-like effector functions, including IL-17A secretion^27^. To further characterize T_reg_17 in this inflammatory setting, we compared frequencies of PD-1^+^, CD69^+^ and CD103^+^ cells within the T_reg_17 and T_reg_ compartment. T_reg_17 cells showed enrichment for these markers relative to total T_reg_ populations in Sham and Abx-treated STNx mice; however, this pattern was not observed in STNx mice (Fig. S3b–d). Additionally, IL-10^+^ T_reg_ were reduced in STNx, but restored under Abx treatment (Fig.S3e), altogether suggesting a loss of suppressive capacity favoring T_H_17-like effector functions in STNx. In line with these findings, STNx induced a sustained T_H_17 and T_reg_17 increase over time (Fig.2i), while T_H_17 of Sham and Abx did not show this response. This parallel expansion of T_reg_17 and T_H_17 cells suggests ineffective regulatory control and persistence of an inflammatory response.

Microbiota-derived metabolites can activate the aryl hydrocarbon receptor (AhR)^28^, which is known to modulate T cell function, including T_H_17 differentiation^29,30^. Consistent with the observed increase in microbiome-derived tryptophan catabolites and predominantly AhR ligand IxS, plasma from STNx mice induced significantly higher AhR activation in a cell-based reporter assay compared with Abx-treated and Sham controls (Fig.2j). Moreover, *Cyp1a1*, a canonical AhR target gene, was upregulated in splenic leukocytes from STNx mice compared to Abx-treated STNx mice (Fig.2k), suggesting immune cells are affected by bacterial AhR ligands.

Next, we evaluated cardiac RORγt+ T cells. We found an increase in T_reg_17, but not T_H_17 cells, in cardiac tissue of STNx mice after 13 weeks (Fig.2l-m, Fig.S3f), consistent with the observation that the cardiac microenvironment promotes T cell transdifferentiation towards regulatory phenotypes^31^. Additionally, infiltrated neutrophils were reduced in STNx mice treated with Abx (Fig.S3g). To further characterize the inflammatory microenvironment of the heart, we performed bulk mRNA sequencing from cardiac tissue. A substantial part of the STNx-associated differentially expressed genes (DEG) were reversed by Abx treatment (Fig.2n). Enrichment analysis on DEG identified Reactome pathways and Gene Ontology (GO) terms involving tissue remodeling (e.g. fibronectin matrix formation, collagen formation, extracellular matrix organization) and immunity (e.g. T cell receptor signaling, interleukin 17 signaling, Fig.2o, Fig.S3h). Variation analysis of IL-17A-associated gene sets revealed a significant increase in enrichment scores in STNx, an effect normalized by Abx (Fig.2p). Taken together, our findings suggest that microbiome-derived, AhR-activating metabolites drive a systemic T_H_17-skewed inflammatory response that impairs T_reg_17 function and promotes adverse cardiac remodeling in kidney disease.

### IL-17A associates with kidney disease severity and adverse outcomes

Given the expansion of IL-17A-producing T cells in experimental CKD, we asked whether a similar inflammatory signature is present in humans and analyzed plasma IL-17A levels in patients with CKD from the UK Biobank (Table S5). Circulating IL-17A concentrations increased progressively with decreasing eGFR and increasing albuminuria, the two main determinants of CKD (Fig.3a, n = 15 120 with available eGFR and albuminuria). IL-17A concentrations were further elevated in patients with eGFR below 60 ml/min/1.73 m^2^ (CKD) and heart failure with preserved ejection fraction (HFpEF, n = 5 497) compared to patients with eGFR below 60 ml/min/1.73 without HFpEF (n = 15 981, Fig.3b). Furthermore, higher plasma IL-17A in individuals with CKD (n = 2 785) was associated with all-cause mortality, even after adjustment for conventional risk factors (Fig.3c-d). These findings substantiate kidney disease-dependent IL-17A response at population scale.

**Figure 3.**
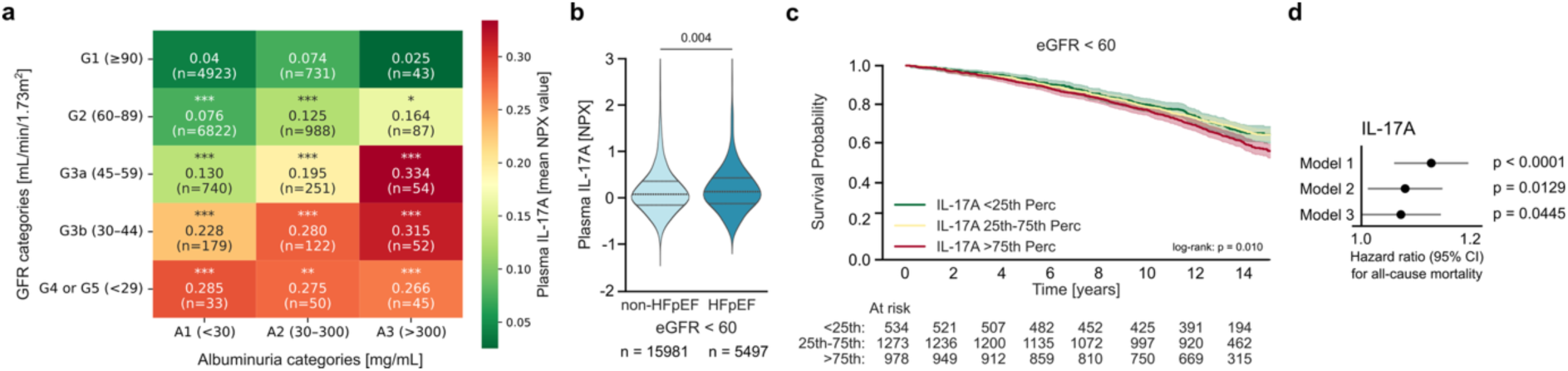
Plasma IL-17A associates with disease severity, cardiovascular comorbidity, and all-cause mortality in patients with CKD. **(a)** Plasma IL-17A derived from OLINK Explore HT proteomics of the UK Biobank (UKBB) stratified by estimated glomerular filtration rate (eGFR) and albuminuria in patients with available data. Statistical comparisons are made between the reference cell (G1, A1) and each corresponding cell. Adjusted p-values indicate significance as follows: *** < 0.001, ** < 0.01, * < 0.05. **(b)** Plasma IL-17A in patients with CKD defined by an eGFR < 60 mL/min/1.73 m². with and without heart failure with persevered ejection fraction (HFpEF). **(c)** Kaplan-Meier survival curves for all-cause mortality stratified by IL-17A levels (<25th, 25th -75th, >75th percentile) among participants with eGFR < 60 mL/min/1.73 m². **(d)** Corresponding hazard ratios and 95% confidence intervals from a Cox proportional hazards model for all-cause mortality. Model 1 only IL-17A. Model 2 additionally adjusted for age, sex, and eGFR. Model 3 further adjusted for hypertension, diabetes, pack years, BMI, and NT-proBNP. **(a-b)** Statistical analyses were performed using one-tailed Student’s t-test or Mann-Whitney U test, as appropriate, followed by Benjamini-Hochberg false discovery rate (FDR) correction for multiple testing. **(c)** Statistical analysis was performed using the log-rank test.

### IxS exacerbates heart failure

IxS has been shown to modulate T_H_17 immunity in neuroinflammation^32^. *In vitro* T_H_17 polarization of murine naïve CD4+ T cells was indeed stimulated dose-dependently by IxS (Fig.4a-b). To test whether IxS stimulates a T_H_17 response *in vivo* and aggravates cardiac remodeling independent of advanced kidney dysfunction, we examined the effects of IxS supplementation in a mouse model of HFpEF^33^ (Fig.4c), the form of heart failure most frequently associated with CKD^3^. We confirmed elevated IxS levels in mice exposed to IxS (Fig.4d), although to a lesser extent compared to STNx mice and patients with CKD^34^, likely due to preserved renal clearance (Fig.4e). Still, IxS levels were sufficient to increase splenic *Cyp1a1* expression (Fig.4f), but not liver *Cyp1a1* (Fig.S4a), indicating that the immune system is particularly susceptible to AhR activation by bacterial metabolites. Similar to STNx mice, splenic IL-17A^+^ T_conv_ (Fig.4g), IL-17A^+^ T_reg_ (Fig.4h) and neutrophils (Fig.S4b) frequencies were elevated in response to IxS. IxS exposure exacerbated heart failure, evidenced by increased heart weights (Fig.4i) and pulmonary congestion (Fig.S4c). Echocardiography confirmed aggravation of diastolic dysfunction (E/e’, E/A, Fig.S4d-e; Table S2) with a preserved left ventricular EF (Fig.S4f; Table S2). Next, we analyzed cardiac fibrosis, the most prominent feature of STNx-associated cardiac remodeling. IxS-exposed hearts displayed higher collagen I and fibronectin content, indicating increased interstitial cardiac fibrosis (Fig.4j-l). Thus, IxS exacerbates heart failure and fibrosis, and elicits a similar inflammatory response as observed in experimental kidney disease.

**Figure 4.**
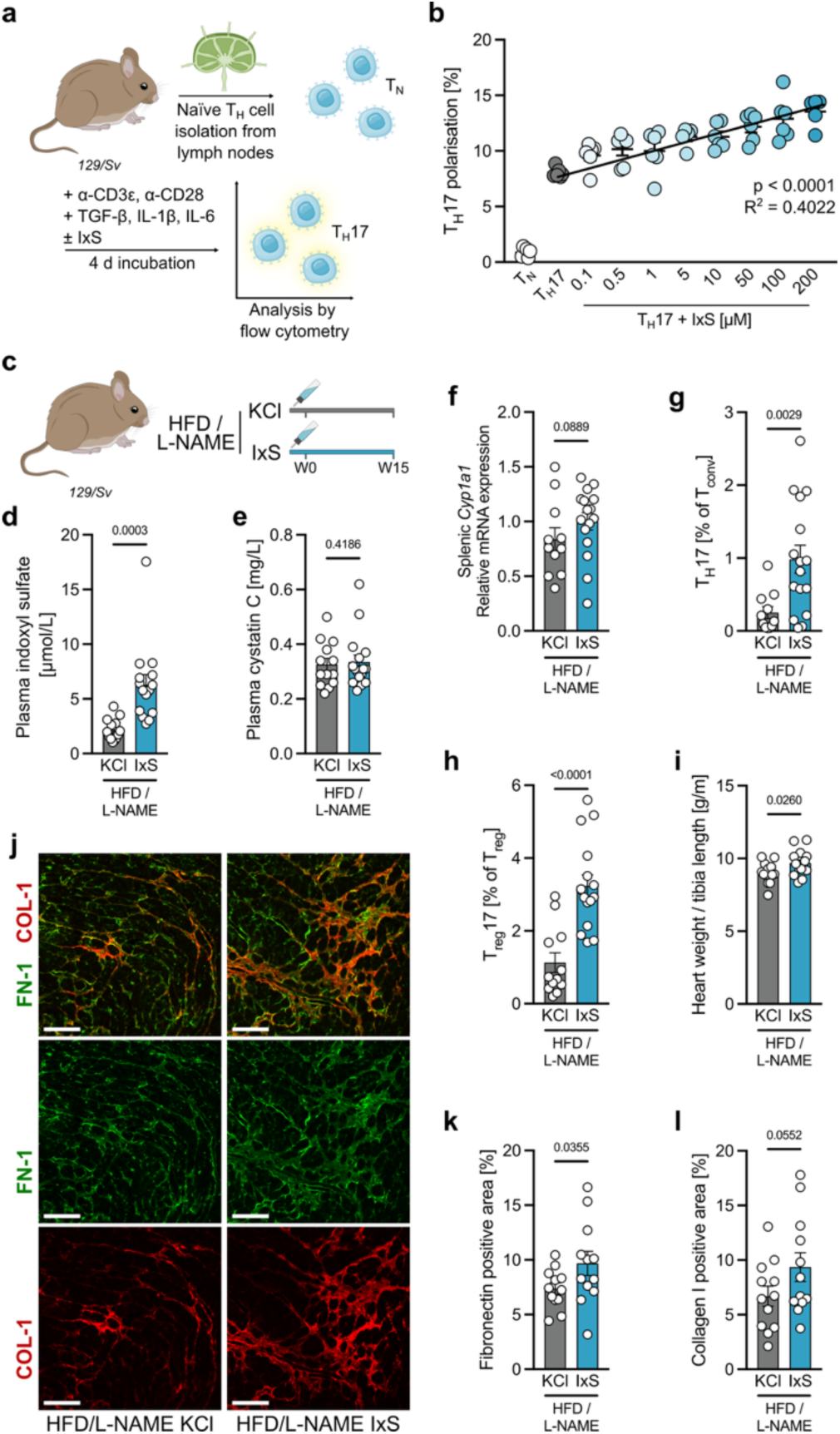
Indoxyl sulfate induces inflammation and cardiac fibrosis independent of kidney function. **(a, b)** *In vitro* T_H_17 differentiation of naïve T helper cells from 129/Sv mice in the presence and absence of indoxyl sulfate (IxS). Two biological replicates, three technical replicates each. **(c)** Heart failure with preserved ejection fraction was induced in 129/Sv mice using high-fat diet (HFD) and L-NAME in the drinking water for 15 weeks. Mice received either IxS or potassium chloride (KCl, control) in the drinking water throughout the experiment. **(d)** Plasma IxS measured by liquid chromatography-mass spectrometry (LC-MS). **(e)** Plasma cystatin C levels. (f) *Cyp1a1* expression in spleen. Flow cytometric quantification of IL17A in splenic **(g)** T_conv_ (CD45^+^ CD3^+^ TCRγδ^−^ CD4⁺ CD8^−^ FoxP3^−^) and **(h)** T_reg_ (CD45^+^ CD3^+^ TCRγδ^−^ CD4⁺ CD8^−^ FoxP3^+^). **(i)** Heart weight normalized to tibia length. **(j)** Cardiac interstitial fibrosis, assessed by histological staining for collagen type I (COL-1) and fibronectin (FN-1) (scale bar: 100 μm), with quantification of **(k)** FN-1 and **(l)** COL-1. Data are shown as mean ± SEM. Each dot represents one mouse. Statistical analyses were performed using one-tailed Student’s t-test or Mann-Whitney U test, as appropriate. **(b)** Statistical analysis was performed using linear regression.

### Cardiac fibroblasts respond to IL-17A and IxS to produce ECM

In addition to the T cell-driven increase in IL-17A production, *Il17ra* expression was upregulated in STNx hearts (Fig.5a), and likewise in IxS-treated HFpEF mice (Fig.5b). To identify IL-17A-responsive cells among cardiac cells, we re-analyzed published single nuclei RNA (sn-RNA) sequencing data from cardiac tissue of STNx and control mice^35^. Among cardiac cell types, fibroblasts showed the highest *Il17ra* expression in STNx (Fig.5c, Fig.S5a). Notably, fibroblasts also showed higher expression of AhR-dependent genes (Fig.S5b) Interestingly, *Il17ra*^+^ fibroblasts displayed higher ECM gene set scores^36^ than *Il17ra*^−^ fibroblasts in STNx mice (Fig.5d). In line, the DEG were characterized by a dominant representation of ECM components (Fig.S5c). To further characterize *Il17ra*^+^ fibroblast transcriptomes, we compared their DEG with different fibroblast signatures known in both mouse and human cardiac tissue^37^. DEG were enriched in myofibroblast-related and, to a lesser degree, cyclic and WNT-related fibroblast-specific genes (Fig.5e), known for their pro-fibrotic and activated status. Cytokine enrichment analysis revealed IL-17A as a prominent factor shaping DEG in cardiac STNx lymphocytes (Fig.S5d). To test the response of cardiac fibroblasts to AhR ligand IxS and IL-17A, respectively, we isolated fibroblasts from healthy murine hearts (Fig.5f). Cell identity and purity were confirmed by immunocytochemistry and flow cytometry (Fig.5g-h). As suggested by the sn-RNA data, IxS upregulated *Il17ra* mRNA expression (Fig.5i), indicating IxS primes fibroblasts to sense IL-17A. Consistent with this, coincubation with IL-17A and IxS resulted in a synergistic increase in *Col1a1*, *Col1a2*, and *Acta2* expression (Fig.5j-l). In line with *in vivo* data, IL17A downregulated *Abi3bp* expression, an ECM protein shown to be reduced in heart failure^38^ (Fig.5m). Immunocytochemistry confirmed the presence of COL1 and IL17RA at the protein level (Fig.5n). Our data demonstrate that IL-17A signaling in cardiac fibroblasts primed by IxS exposure leads to increased ECM production, ultimately exacerbating cardiac fibrosis.

**Figure 5.**
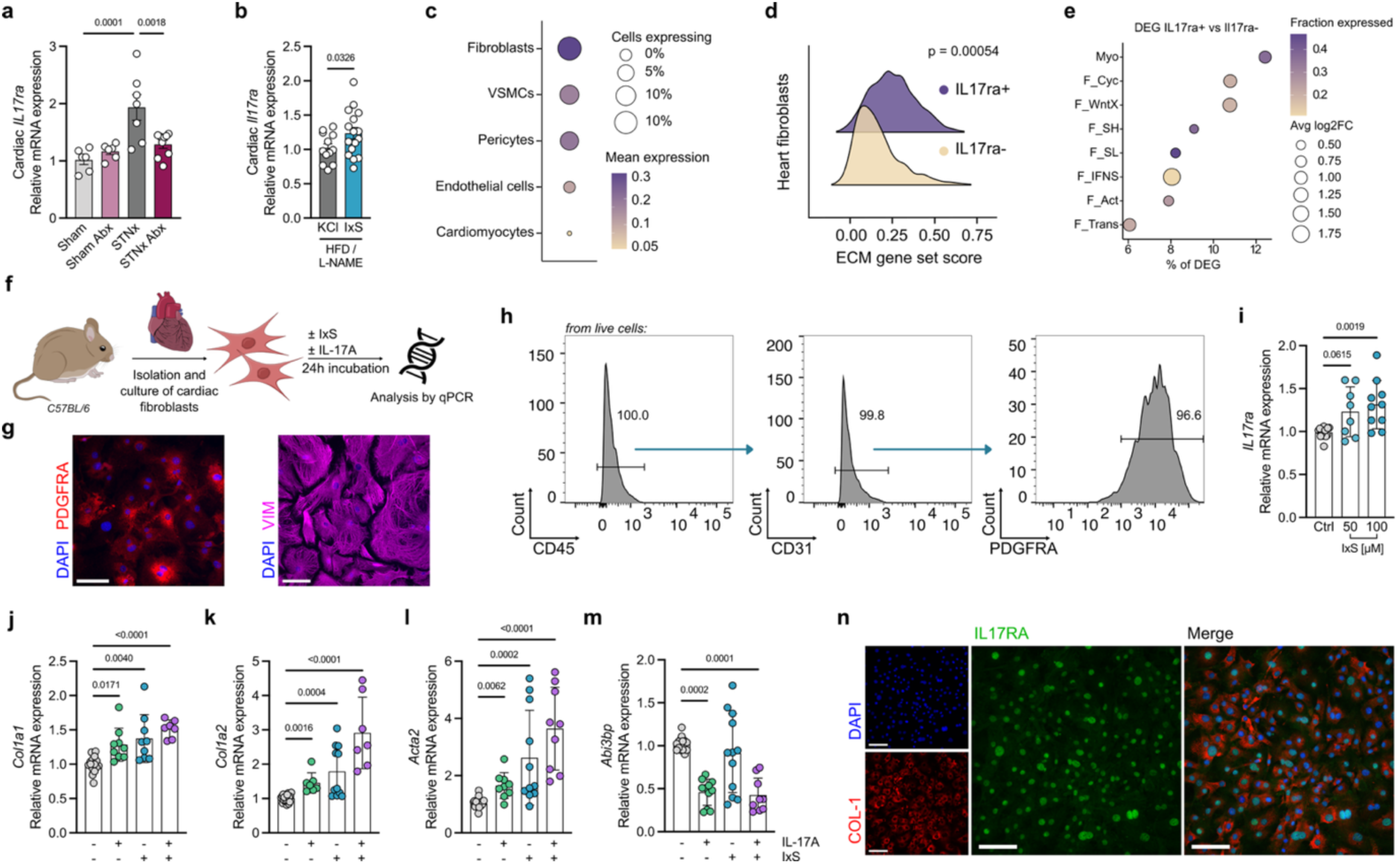
IL-17A-signaling in IxS-primed cardiac fibroblasts promotes extracellular matrix production. Cardiac *Il17ra* expression in **(a)** 129/Sv mice subjected to subtotal nephrectomy (STNx) or sham operation and treated with either an antibiotic cocktail (Abx) or control drinking water, and in **(b)** 129/Sv mice subjected to high-fat diet (HFD) plus L-NAME and indoxyl sulfate (IxS) or potassium chloride (KCl, control) in the drinking water. Re-analysis of previously published cardiac single nuclei (sn) RNA-sequencing data (Kaesler *et al.*) from STNx mice shows **(c)** highest *Il17ra* expression in fibroblasts and **(d)** higher extracellular matrix (ECM) gene set score in *Il17ra* expressing cardiac fibroblasts. **(e)** Enrichment analysis for known cardiac fibroblasts subtypes within differentially expressed genes. **(f)** Cardiac fibroblasts were isolated from C57BL/6J mice for *ex vivo* treatment with IxS and IL-17A. Purity validation **(g)** by histology using platelet-derived growth factor receptor alpha (PDGFRA) and vimentin (VIM) (scale bar: 100 μm), and **(h)** by flow cytometry showing CD45^−^ CD31^−^ PDGFRA^+^ fibroblasts. **(i)** *Il17ra* expression following *in vitro* incubation with IxS. Isolated fibroblasts were incubated with 100µM IxS, 1ng/mL IL-17A, or both and expression of **(j)** *Col1a1*, **(k)** *Col1a2*, **(l)** *Acta2*, **(m)** *Abi3bp* were quantified. **(n)** Histological confirmation of IL-17RA expression in cardiac fibroblasts (scale bar: 100 μm). **(a, b)** Data are shown as mean ± SEM. Each dot represents one mouse. **(i-m)** Data are combined from four biological replicates, with three technical replicates each. Statistical analyses were performed using one-way ANOVA or Kruskal-Wallis test followed by Tukey’s or Dunn’s *post hoc* correction, as appropriate. **(b)** Statistical analysis was performed using one-tailed Student’s t-test. **(d)** Comparison by two-sided Mann-Whitney U-test.

### AhR inhibition ameliorates cardiac remodeling in kidney disease

Since STNx-associated cardiac remodeling occurs in an AhR-activating milieu that in turn promotes IL-17A production and cardiac fibrosis, we investigated whether AhR inhibition confers therapeutic benefit. For optimal translational potential, we re-purposed the orally bioavailable AhR inhibitor BAY 2416964 (AhRi), currently under investigation in solid malignancies^39^. *In vitro*, AhRi dose-dependently reduced the IxS-induced upregulation of *Cyp1a1* expression in murine splenocytes (Fig.6a-b). Furthermore, AhRi dose-dependently suppressed IxS-driven T_H_17 polarization (Fig.6a,c). In both assays, AhRi was effective at doses of 10^−7.5^-10^−6.5^ M. Next, we aimed to test the efficacy of AhRi *in vivo* in STNx mice (Fig.6d). Therefore, mice were stratified according to their CysC levels (Fig.S6a) and randomized to receive either AhRi or vehicle starting two weeks after complete STNx through to week 13 by daily oral gavage. Sham-operated vehicle-treated mice served as additional control. At the endpoint, we detected similar levels of CysC in AhRi- and vehicle-treated mice (Fig.S6b). AhRi treatment improved survival of STNx mice (Fig.6e) and lowered systemic inflammation, as measured by plasma CRP (Fig.6f). Upregulation of canonical AhR target gene *Cyp1a1* in immune cells was prevented by AhRi (Fig.6g), as well as the STNx-associated increase in both T_H_17 (Fig.6h) and T_reg_17(Fig.6i). In line, IL-17A+ T_conv_ and T_reg_ were reduced by AhRi (Fig.S6c-d). We observed an improvement of the cardiac phenotype upon AhRi reminiscent of the Abx effect, with lower left ventricular mass (Fig.6j), and improved cardiac function (Fig.6k-l, Table S3) and reduced fibrosis (Fig.6m-n), together with lower cardiac *Il17ra* expression (Fig.6o). Bulk transcriptomic analyzes of cardiac tissue showed similar effects as observed with Abx treatment, with a reduction in GO terms and pathways involved in ECM formation and inflammatory processes (Fig.6p, Fig.S6e). Targeted analysis of T_H_17-related signaling pathways revealed significantly higher scores upon STNx, which were rescued by AhRi (Fig.6q). These findings identify the AhR as a key mediator of inflammation and cardiac remodeling in kidney disease and suggest AhRi as a therapeutic strategy.

**Figure 6.**
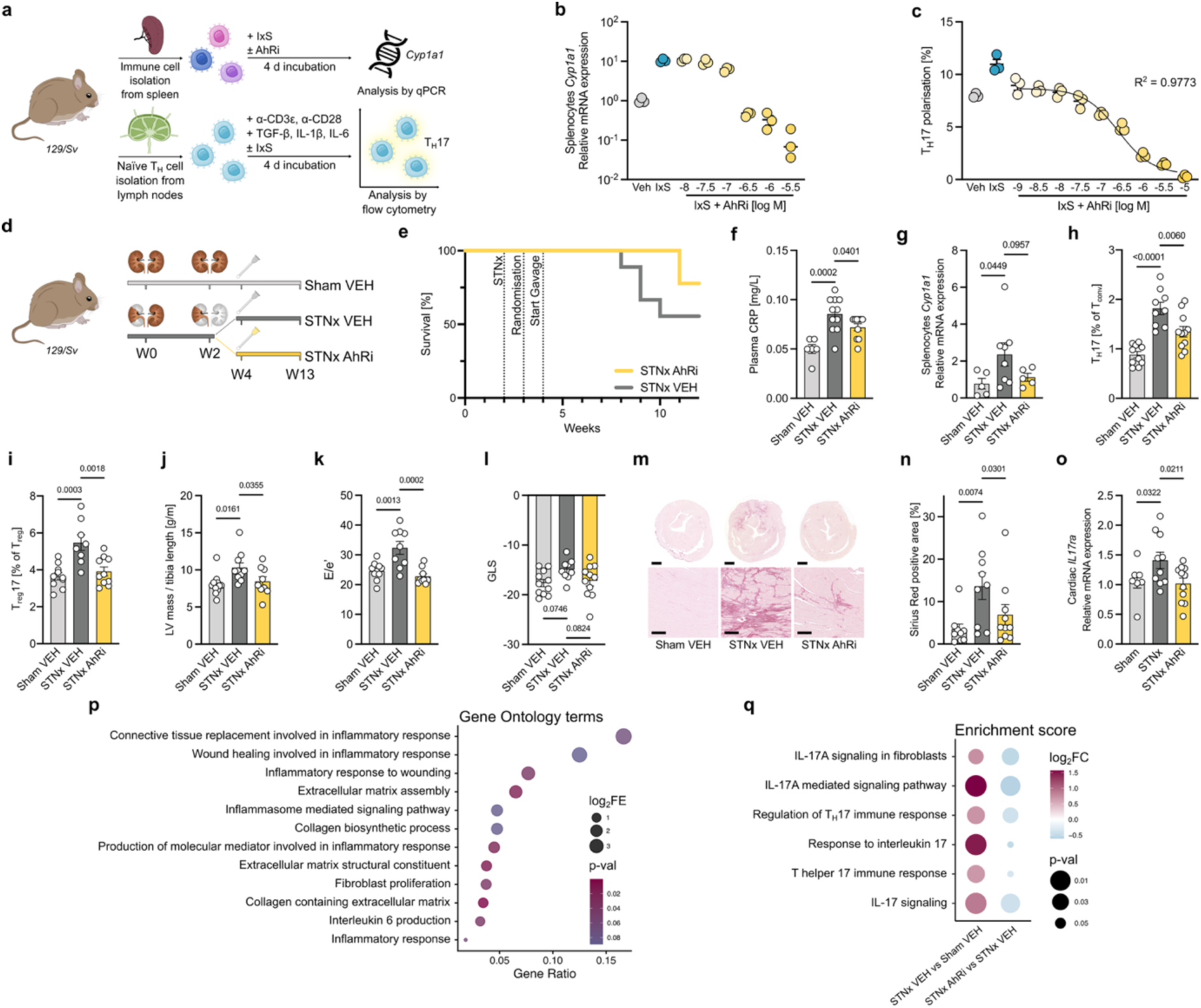
Pharmacological blockade of the AhR ameliorates inflammation and cardiac remodeling in kidney disease. **(a)** *Ex vivo* aryl hydrocarbon receptor (AhR) inhibition in immune cells from 129/Sv mice with BAY 2416964 (AhRi). **(b)** AhRi blocks indoxyl sulfate (IxS)-induced *Cyp1a1* gene expression in splenocytes. **(c)** AhRi suppresses IxS-enhanced TH17 differentiation of naïve T cells. **(d)** 129/Sv mice underwent two stage subtotal nephrectomy (STNx) or sham surgery. Two weeks after completion of surgery, STNx mice were randomized to AhRi or vehicle (VEH) treatment based on plasma cystatin C. **(e)** Survival analysis according to pre-specified humane endpoints. **(f)** Plasma C-reactive protein (CRP) levels. **(g)** *Cyp1a1* expression in splenocytes. RORγt in splenic **(h)** T_conv_ (CD45^+^ CD3^+^ TCRγδ^−^ CD4⁺ CD8^−^ FoxP3^−^) and **(i)** T_reg_ (CD45^+^ CD3^+^ TCRγδ^−^ CD4⁺ CD8^−^ FoxP3^+^) by flow cytometry. **(j)** Left ventricular (LV) mass normalized to tibia length, **(i)** E/e’ ratio and **(l)** global longitudinal strain (GLS) from endpoint echocardiography. **(m)** Cardiac fibrosis assessed by Sirius Red staining (scale bar top: 1mm, bottom: 100µm). **(n)** Quantification of Sirius Red-positive interstitial fibrosis. **(o)** Cardiac *Il17ra* expression. **(p)** Enriched biological pathways among differentially expressed genes significant in STNx rescued by AhRi. **(q)** Targeted gene ontology (GO) analysis of IL-17A-related differentially expressed gene sets. **(b-c)** Data are combined from three technical replicates. **(c)** Data were analyzed by nonlinear sigmoidal regression. **(e-o)** Data are shown as mean ± SEM. Each dot represents one mouse. Statistical analyses were performed using one-way ANOVA or Kruskal-Wallis test followed by Tukey’s or Dunn’s *post hoc* correction, as appropriate, unless otherwise indicated. Gene set enrichment analyses were performed using Fisher’s exact test.

### Conserved pro-fibrotic IxS-AhR-T_H_17 axis in humans

We further aimed to explore the translational potential of AhRi by investigating the efficacy in human cells. AhRi dose-dependently blocked the IxS-stimulated AhR activity in human AhR reporter cells (Fig.7a). Likewise, AhRi reduced the AhR activating potential of serum collected from patients with kidney failure requiring hemodialysis (HD) to levels observed with serum from healthy controls (HC, Fig.7b-c; Table S4). Moreover, addition of HD serum to human T_H_17 polarization assays increased T_H_17 cells compared to HC (Fig.7b,d). Hence, the kidney disease-responsive AhR-T_H_17 axis identified in mice appears to be conserved in humans.

**Figure 7.**
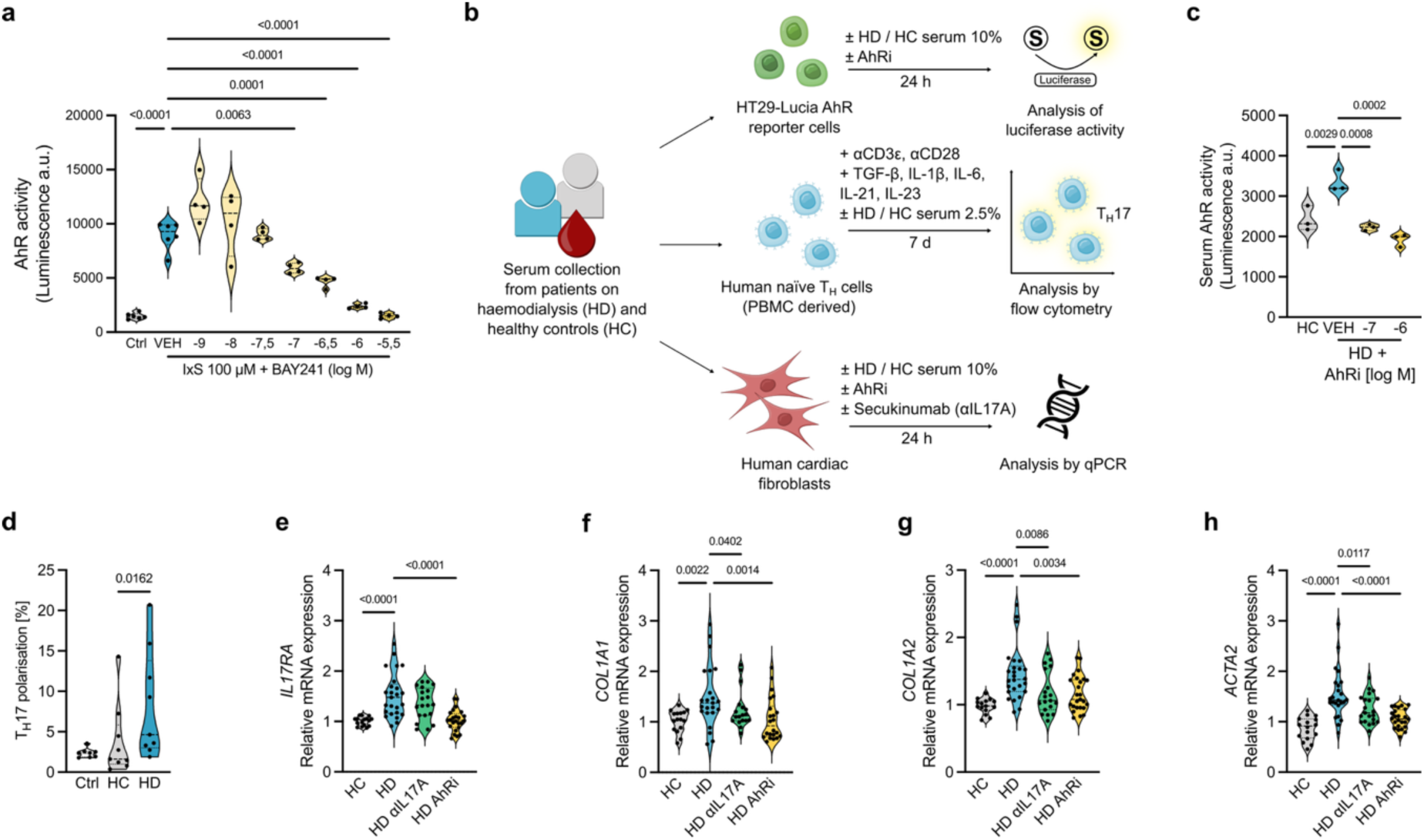
Serum from patients with CKD increases T_H_17 differentiation and cardiac fibroblast activation. **(a)** Activation of aryl hydrocarbon receptor (AhR) in HT29-Lucia AhR reporter cells with 100µM indoxyl sulfate (IxS) is blockable by the AhR inhibitor BAY 2416964 (AhRi). **(b)** Serum was obtained from healthy controls (HC) or patients with chronic kidney disease treated with hemodialysis (HD) for *in vitro* testing on AhR reporter cells, naïve T cells, and human cardiac fibroblasts. **(c)** Activation of the AhR in response to HC or HD serum in the presence or absence of AhRi. **(d)** *In vitro* T_H_17 differentiation of naïve T cells from healthy volunteers in the presence of HC and HD serum. **(e-h)** Human primary cardiac fibroblasts were incubated *in vitro* with HC and HD serum in addition of anti-IL-17A (Secukinumab, αIL17A) or AhRi and analyzed for **(e)** *IL17RA*, **(f)** *COL1A1*, **(g)** *COL1A2*, and **(h)** *ACTA2* expression. **(a, c, d)** Data are combined from three biological replicates, with three technical replicates each and **(e-h)** data combined from two primary cell lines from different donors treated with HC serum (n = 4 donors) or HD serum (n = 9 donors) in triplicates. Statistical analyses were performed using one-way ANOVA or Kruskal-Wallis test followed by Tukey’s or Dunn’s *post hoc* correction, as appropriate.

To verify the role of AhR signaling and IL-17A in fibroblast activation in humans, we used publicly available single cell RNA (sc-RNA) sequencing data. Since cardiac sc-RNA data from patients with CKD were not available, we explored data from kidney tissue of patients with CKD and healthy controls^40^. Fibroblasts of patients with CKD displayed the highest AhR enrichment scores compared to other kidney cell types (Fig.S7a). Moreover, myofibroblasts expressed the highest levels of *IL17RA* (Fig.S7b), and *IL17RA* expressing myofibroblasts exhibited markedly higher ECM module scores (Fig.S7c). In line, pro-fibrotic and ECM-associated genes were among the major DEG (Fig.S7d), indicating that IL-17A-responsive fibroblasts represent a pro-fibrotic subset.

Next, we stimulated primary human cardiac fibroblasts with serum from hemodialysis patients (HD) or healthy controls (HC) (Fig.7b). HD serum induced upregulation of *IL17RA* (Fig.7e, Fig.S8a) together with ECM-associated transcripts such as *COL1A2* and *ACTA2* (Fig.7f-h, Fig.S8b-d). We tested whether these effects could be prevented by AhRi or secukinumab, a human monoclonal antibody used in clinical routine targeting IL-17A (αIL-17A). Both AhRi and αIL-17A mitigated the pro-fibrotic effects (Fig.7f-h, Fig.S8b-d). However, only AhRi reduced *IL-17RA* expression (Fig.7e, Fig.S8a). Similar to murine cardiac fibroblasts, IxS upregulated *IL17RA* expression in primary human cardiac fibroblasts, an effect abrogated by AhRi (Fig.S8e). Both IxS and IL-17A increased *COL1A2* and *ACTA2* expression, whereas co-stimulation with both ligands again had a synergistic effect, leading to the highest pro-fibrotic transcriptional response (Fig.S8f-h). Proliferation was not altered with IL-17A and IxS co-incubation, as determined by Ki-67 staining (Fig.S8i-j). Taken together, interrogating data from human sc-RNA sequencing and *in vitro* assays using patient-derived serum show a conserved pro-fibrotic IxS-AhR-T_H_17 axis in CKD.

## Discussion

This study demonstrates that kidney disease recalibrates microbiome-host interactions to potentiate inflammation-mediated cardiac fibrosis. We identify a microbial AhR-activating metabolite driving a pathogenic T_H_17 response that engages IL-17A-responsive cardiac fibroblasts, positioning the metabolite-AhR-IL-17A axis as a therapeutic target to prevent cardiac fibrosis in kidney disease.

Inflammation is a key driver of CVD. In CKD, inflammation is substantially elevated, paralleling a markedly increased risk for CVD^41^. Evidence suggests that CVD development in CKD extends beyond traditional risk factors^42^ such as hypertension, hyperlipidemia, and diabetes, yet underlying mechanisms remain less characterized. Adverse cardiac remodeling and fibrosis are prevalent in CKD^43^, with heart failure representing a frequent clinical manifestation^44^. The microbiome plays a central role in immune modulation, in part via action of bacterial metabolites. In CKD, these metabolites accumulate and contribute to a pro-inflammatory internal milieu. Such metabolites were shown to be of bacterial colonic origin and can reach toxic levels in CKD^8^. To date, therapeutic strategies aimed at improving outcomes by reducing bacterial metabolite levels through unspecific intestinal adsorption showed limited success^45^, underscoring the need for a refined understanding of microbiome-host interaction in CKD.

Our mechanistic investigations were prompted by the observation that the depletion of the gut microbiome in a state of reduced kidney function drastically reduced the development of cardiac fibrosis as well as diastolic dysfunction, and suppressed systemic inflammation, as evidenced by a lower plasma CRP. Remarkably, cardioprotection occurred despite persistently high blood pressure, in line with a clinical observation showing that cardiac fibrosis in CKD occurs independent of hypertension^43^, which has so far not been linked to the gut microbiome. The profound inflammation encouraged us to investigate cellular immunity further, in which a microbiome-dependent T_H_17 response to STNx became evident. While IL-17A has previously been associated to ischemic heart disease^5^ and kidney disease progression after acute kidney injury^46^, it has not been highlighted as a cytokine linking heart disease and CKD. Our analyzes of UK Biobank data corroborate this link, demonstrating elevated IL-17A levels in patients with CKD and HFpEF. IL-17A plasma concentrations increased with increasing severity of CKD, reflected by decreasing eGFR or increasing albuminuria and were associated with excess mortality.

We previously hypothesized that CKD-linked inflammation is mediated by the AhR^30^. Plasma AhR activation potential and AhR-dependent gene expression in splenocytes increased following STNx; both effects being abrogated by microbiome depletion. We also present evidence that AhR activation and T_H_17 response are mediated via the AhR by increased levels of IxS, a known microbiome-derived AhR-activating ligand. *In vitro*, IxS induced T_H_17 polarization dose-dependently. *In vivo*, oral administration of IxS in an experimental heart failure model increased AhR activation and AhR-dependent gene expression in immune cells, promoted T_H_17 expansion, and aggravated cardiac fibrosis, despite preserved kidney function. Further *in vivo* evidence is provided by AhR inhibition in STNx mice, which abrogated AhR-dependent gene expression in immune cells, T_H_17 expansion and systemic inflammation, as well as cardiac fibrosis and diastolic dysfunction. Notably, AhR inhibition conferred cardioprotection without improvement in kidney function, consistent with prior evidence that IL-17A is dispensable for renal pathology in experimental CKD^47^. Thus, IxS-dependent AhR activation appears to be central to cardiac remodeling in kidney disease.

As an environmental sensor integrating various external and endogenous cues, the AhR is known to regulate T_H_17 differentiation^48^. AhR-deficient naïve T cells exhibit reduced differentiation^48^, while AhR activation by high affinity ligands has been shown to promote T_H_17 and autoimmune disease^49^. However, AhR-mediated effects on T_H_17 differentiation depend on the ligand and cytokine milieu^50^. IxS is a known endogenous AhR ligand and has been linked with inflammatory milieus as well as T_H_17-mediated autoimmunity^32^. We present evidence that IxS activates AhR transcriptional programs in CD4+ cells to promote a pathogenic T_H_17 response in kidney disease. We show that IxS drives RORγt expression and IL-17A production in CD4+ T cells *in vitro* and *in vivo*. Intriguingly, we also detected increased IL-17A production by Foxp3+ CD4+ T_reg_, as well as increased RORγt expression in CD4+ FoxP3+ cells (T_reg_17), especially in cardiac tissue. While T_reg_17 cells typically restrain IL-17A-driven inflammation through IL-10^51^, their regulatory capacity appears diminished in the present uremic setting, where IL-17A-mediated inflammation is less effectively controlled. Consistently, T_reg_17 cells from uremic STNx mice exhibited reduced expression of PD-1, CD103, and CD69, proteins implicated in the maintenance of suppressive function^32^, compared to microbiome-depleted mice with kidney failure and healthy control mice. This phenotype suggests a functional shift in T_reg_17 cells toward RORγt-driven effector activity at the expense of FoxP3-dependent regulatory functions. Moreover, IL-10 production was lowest in T_reg_ from STNx mice. The STNx-induced IL-17A response was accompanied by neutrophilia and enhanced cardiac neutrophil infiltration. IL-17A is a known regulator of neutrophils, promoting granulopoiesis and inducing neutrophil-attracting chemokines in stromal and cardiac cells^52^. Furthermore, circulating neutrophils correlate with IL-17A levels^25^. Once recruited, neutrophils maintain IL-17A-driven inflammation through release of inflammatory mediators, including IL-1β and neutrophil extracellular traps^53^. Together, these findings reveal a microbiome-dependent, AhR-mediated and IL-17A-driven inflammatory program in kidney disease.

Our study identifies cardiac fibroblasts as key targets of IL-17A, characterized by enhanced IL-17RA expression *in vivo*, which in turn drives ECM production and promotes cardiac fibrosis. It has previously been reported that IL-17A enhances fibroblast proliferation and migration^54^ and induces a proinflammatory fibroblast phenotype involving GM-CSF^55^. We demonstrate that IxS effectively primes cardiac fibroblasts via AhR-dependent upregulation of IL-17RA in murine and human cells, suggesting that the same bacterial metabolite that drives IL-17A-mediated inflammation also sensitizes fibroblasts to IL-17A signaling and ECM production. Thus, IxS orchestrates inflammation and tissue remodeling through AhR signaling, providing a mechanistic explanation for the pronounced cardiac fibrosis observed in CKD and a strong rationale for AhR inhibition as a therapeutic strategy. Notably, pharmacological AhR targeting with a small-molecule inhibitor currently under clinical investigation for cancer treatment^39^ (NCT04999202) proved effective both *in vivo* in STNx mice and *in vitro* in human cardiac fibroblasts, highlighting its translational potential. Targeting host sensing of uremic toxins may uncouple disease-driving microbiome-derived metabolites from downstream inflammatory responses. Integrating AhR inhibition with existing therapies could therefore provide a complementary strategy to mitigate adverse cardiovascular comorbidities in CKD.

Taken together, our findings identify an IxS-AhR-IL-17A axis as a central driver of cardiac inflammation and fibrosis in kidney disease. We thereby demonstrate how kidney dysfunction promotes cardiac pathology through specific microbiome-derived metabolites. Our study positions IxS not merely as a biomarker of uremia but as an active pathogenic effector linking microbial metabolite accumulation to maladaptive immunity and tissue fibrosis via AhR signaling. AhR inhibition may offer a means to selectively target uremic downstream signaling while preserving potentially beneficial microbiome functions. More broadly, these findings refine our knowledge of disease-to-multimorbidity transition, moving beyond traditional organ-centered models toward a mechanistic understanding of inter-organ communication and highlighting the microbiome as an upstream driver of host response reprogramming that facilitates pathological organ crosstalk.

## Author contribution

CRediT: Conceptualization: AY, WA, AT, JH, HB, NW; Data curation: AY, WA, JVe, HA, MK; Formal Analysis: AY, WA, OP; Funding acquisition: VM, UK, AZ, PO, WG, KUE, DNM, MK, HB, NW; Investigation: AY, WA, AT, OP, FS, IAS, AMR, JVe, NG, VM, SH, SK, GT, HA, FF, CH, SAH, PMB, MIW, FB; Methodology: AY, WA, OP, FS, IAS, JVe, JVo, PO, WG, KD, MK, HB, NW; Project administration: AY, WA; Resources: UK, TK, BK, AZ, PO, WG, KD, KUE, DNM, MK, JH, HB, NW; Software: AY, WA, JVe, HA; Supervision: JVo, UK, TK, BK, AZ, PO, WG, KD, KUE, DNM, MK, JH, HB, NW; Validation: AY, WA, RK, MK; Visualization: AY, WA, JVe, HB, NW; Writing - original draft: AY, WA, HB, NW; Writing - review & editing: all authors.

## Acknowledgments

We thank Melanie Röhr, Gudrun Koch, Laura Mossier, and Daniel Herrmann for their excellent technical assistance, Ilona Kamer for the implantation of the blood pressure telemetry devices and Heike Schwede for support with animal husbandry. Clinical chemistry and echocardiography measurements were performed at the Preclinical Research Center of the MDC. Bulk mRNA-seq was performed at the BIH/MDC Genomics Technology Platform. Human T cells were sorted using the flow cytometry platform of the MDC. We thank the MDC Max Cluster for providing us with the scientific compute cluster resources and Ulrike Löber for bioinformatics support.

