## Supplementary Figures S1-S9, Supplementary Tables S1-S8, Supplementary Methods for "Kidney disease reprograms microbiome-host signaling to promote heart failure"

Table of content

| Supplementary figures | |
|---|---|
| Figure S1 | Characterization of gut microbiota in mice with impaired kidney function |
| Figure S2 | Depletion of microbiome-derived metabolites influence cardiorenal damage in mice with impaired kidney function |
| Figure S3 | Kidney disease-driven inflammation is ameliorated in gut microbiome-depleted mice |
| Figure S4 | Indoxyl sulfate exacerbates inflammation and cardiac injury in a mouse model of diastolic heart failure |
| Figure S5 | Phenotype of *Il17ra* expressing cardiac fibroblasts |
| Figure S6 | Pharmacological blockade of the aryl hydrocarbon receptor ameliorates kidney disease-driven inflammation |
| Figure S7 | Phenotype of *IL17RA* expressing human fibroblasts in kidney tissue from patients with chronic kidney disease |
| Figure S8 | Pro-fibrotic effects of indoxyl sulfate and IL-17A on human primary cardiac fibroblasts |
| Figure S9A | Gating strategy for mouse flow cytometry analysis of panel 1 and panel 2 |
| Figure S9B | Gating strategy for mouse flow cytometry analysis of panel 3 and panel 4 |
| Supplementary tables | |
| Table S1 | Echocardiographic parameters of the Abx trial |
| Table S2 | Echocardiographic parameters of the IxS trial |
| Table S3 | Echocardiographic parameters of the AhRi trial |
| Table S4 | Baseline characteristics of serum donors CKD stage 5D and healthy controls |
| Table S5 | Baseline characteristics of the UK Biobank participants stratified by eGFR |
| Table S6 | Overview of primers and probes used in this study |
| Table S7 | Overview of mouse antibodies used in this study |
| Table S8 | Overview of human antibodies used in this study |
| Supplementary methods | |

### Supplementary figures

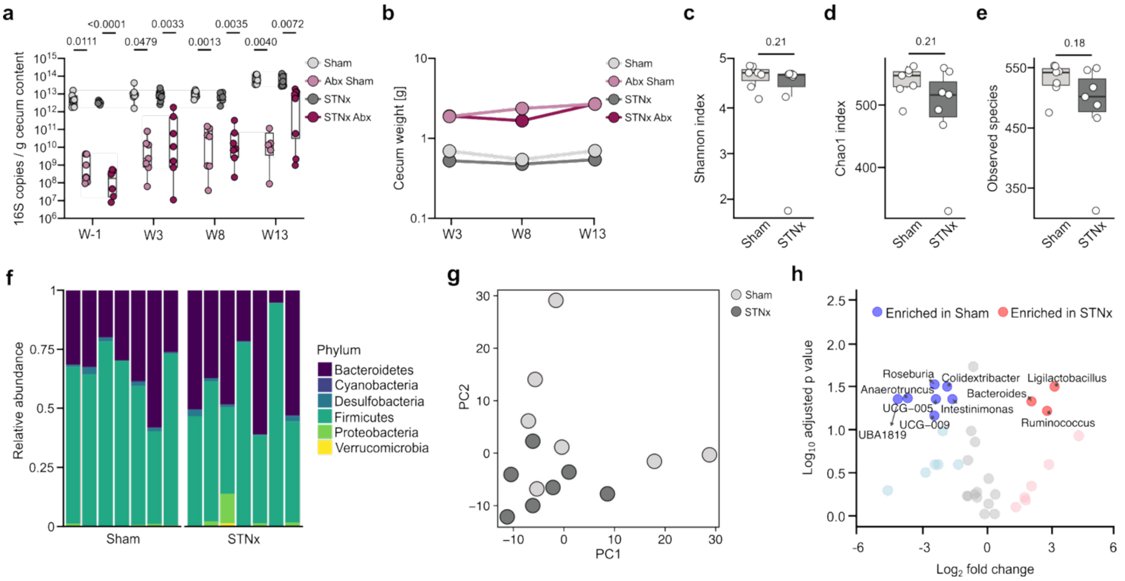

**Figure S1. Characterization of gut microbiota in mice with impaired kidney function.** 129/Sv mice were randomized to receive a broad-spectrum, non-absorbable oral antibiotic cocktail (Abx) or control treatment. After one week, mice underwent two-stage subtotal nephrectomy (STNx) or sham surgery. Organs were collected at given time points: week (w) -1, w3, w8, w13. **(a)** 16S rDNA copies per g of cecal content quantified by qPCR. **(b)** Cecum weights over the course of the experiments. 16S rDNA sequencing was performed at week 13 and alpha ($\alpha$) diversity was analyzed by **(c)** Shannon index, **(d)** Chao1 index, and **(e)** observed amplicon sequencing variants (ASV). **(f)** Relative abundance of phyla, each bar represents one individual animal. **(g)** Principal component analysis (PCA) of ASV-level microbiome data. **(h)** Differential abundance analysis on genus level. Differences were assessed using two-sided Mann-Whitney U test with false discovery rate correction according to Benjamini-Hochberg for **(a)** and **(h)**; taxa were considered differentially abundant at FDR <0.1 and $|\log_2$ fold change$| > 1$.

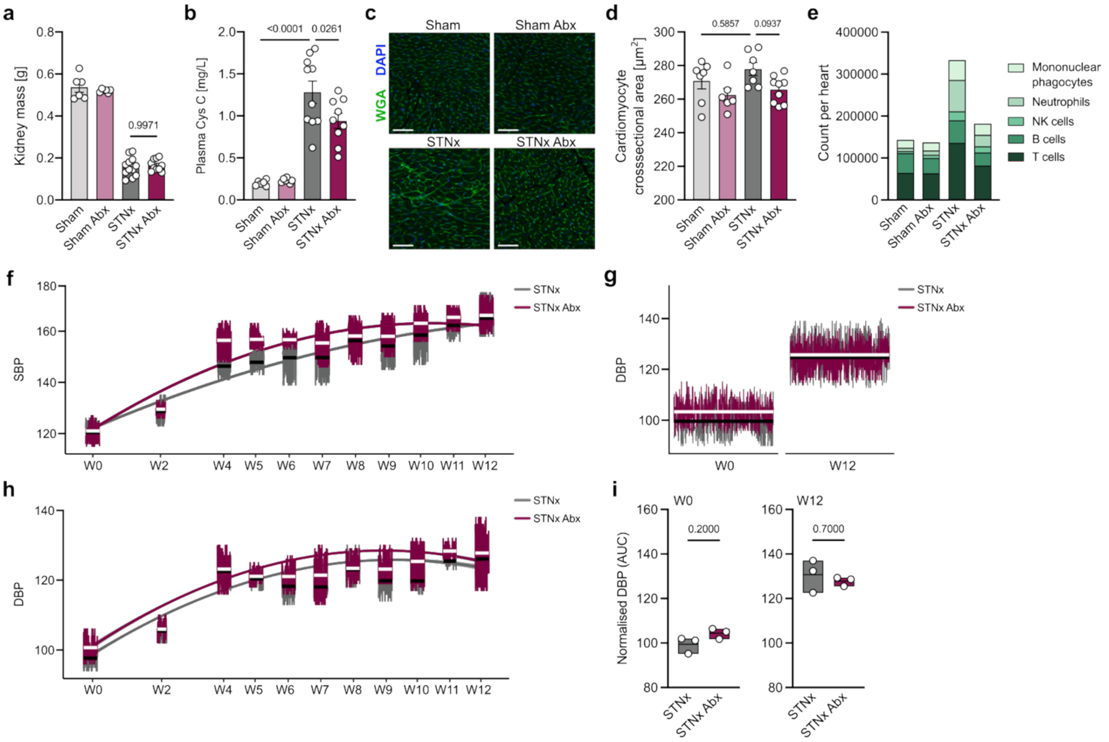

**Figure S2. Depletion of microbiome-derived metabolites influences cardiorenal damage in mice with impaired kidney function. (a)** Residual kidney mass in 129/Sv mice after two stage subtotal nephrectomy (STNx) or Sham operation with concomitant treatment with an oral antibiotic cocktail (Abx) or control. **(b)** Plasma cystatin C (Cys C) levels. **(c)** Histological assessment of cardiomyocyte hypertrophy by wheat germ agglutinin (WGA) staining (scale bar: 50 µm). **(d)** Quantification of cardiomyocyte cross-sectional area. **(e)** Absolute numbers of leucocyte populations infiltrating cardiac tissue. **(f)** Radiotelemetry analysis of systolic blood pressure (SBP) over the course of the experiment. **(g)** Average diastolic blood pressure (DBP) at baseline (week 0) and week 12. **(h)** Radiotelemetry analysis of DBP over the course of the experiment. **(i)** Individual mean DBP at week 0 and week 12. Statistical analyses were performed using one-way ANOVA or Kruskal-Wallis test followed by Tukey's or Dunn's *post hoc* correction, as appropriate, unless otherwise indicated. Two-group comparisons of blood pressure in **(i)** were assessed using Mann-Whitney U test.

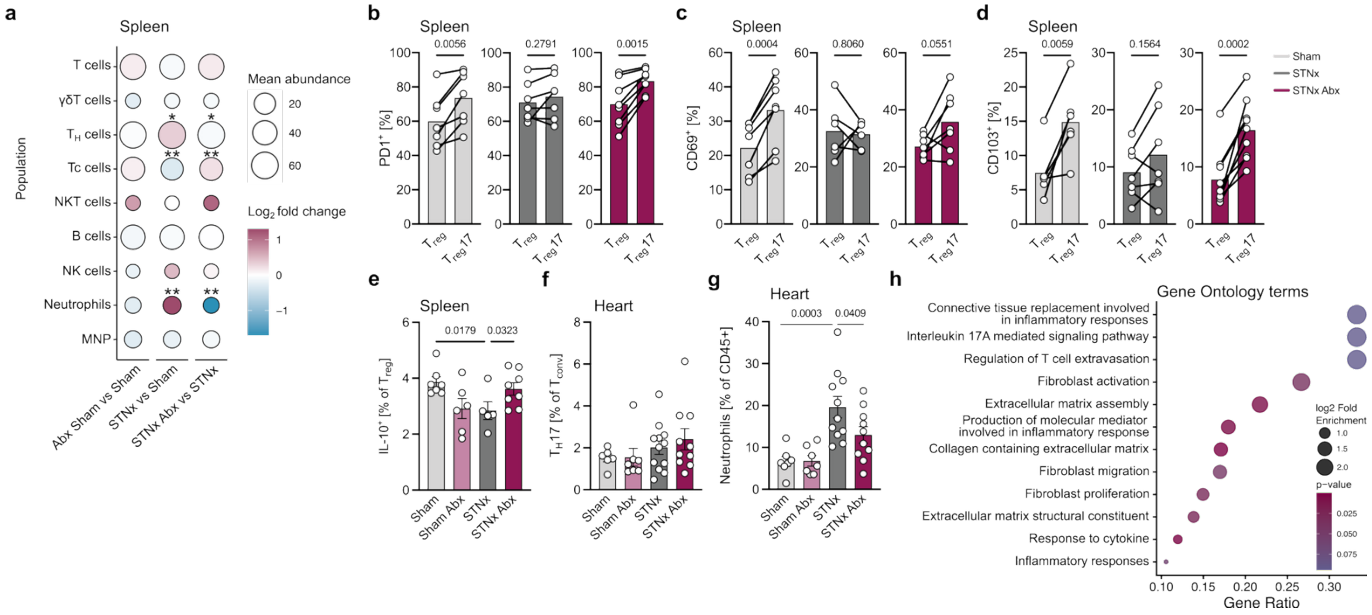

**Figure S3. Kidney disease-driven inflammation is ameliorated in gut microbiome-depleted mice.** 129/Sv mice received two stage subtotal nephrectomy (STNx) or Sham operation with concomitant treatment with an oral antibiotic cocktail (Abx) or control. **(a)** Comparison of major splenic immune populations analyzed by flow cytometry (* p-adj. < 0.1, ** p-adj. < 0.05). Comparison of splenic $T_{reg}$ ($CD45^+$ $CD3^+$ $TCR\gamma\delta^-$ $CD4^+$ $CD8^-$ $Foxp3^+$) and $T_{reg}17$ ($ROR\gamma t^+$ $T_{reg}$) for **(b)** PD-1, **(c)** CD69, and **(d)** CD103 expression. **(e)** IL-10 expression in splenic Treg. **(f)** RORγt expression in cardiac $T_{conv}$ ($CD45^+$ $CD3^+$ $TCR\gamma\delta^-$ $CD4^+$ $CD8^-$ $Foxp3^-$). **(g)** Relative proportion of neutrophils ($CD45^+$ $CD3^-$ $CD19^-$ $NK1.1^-$ $CD11b^+$ $Ly6G^+$) among leucocytes isolated from the heart. **(h)** Significantly enriched gene ontology (GO) terms in the cardiac transcriptome. Data are shown as mean ± SEM. Each dot represents one mouse. Statistical testing was performed using Benjamin-Hochberg with FDR for the bubble plot, paired *t* test for the two-group comparison **(b-d)** and one-way ANOVA followed by Tukey's *post hoc* correction for the four-group comparison. Gene set statistical analyses were performed using Fisher's exact test followed by FDR-correction.

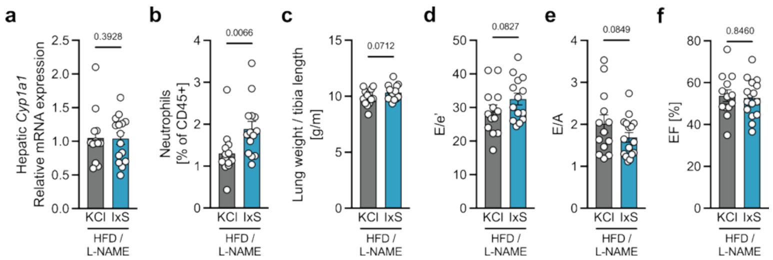

**Figure S4. Indoxyl sulfate exacerbates inflammation and cardiac injury in a mouse model of heart failure.** Heart failure with preserved ejection fraction was induced in 129/Sv mice using high-fat diet (HFD) and L-NAME in the drinking water for 15 weeks. Animals received either IxS or potassium chloride (KCl) as control in the drinking water over the course of the experiment. **(a)** *Cyp1a1* expression in the liver. **(b)** Relative abundance of neutrophils ($CD45^+$ $CD3^-$ $CD19^-$ $NK1.1^-$ $CD11b^+$ $Ly6G^+$) in the spleen. **(c)** Lung weight normalized to tibia length. Echocardiographic assessment of **(d)** E/e’, **(e)** E/A, and **(f)** ejection fraction (EF). Data are shown as mean ± SEM. Each dot represents one mouse. For confirmatory analysis, one-tailed Student’s t test or Mann-Whitney U test was used, as appropriate. In **(f)**, statistical analysis was performed using two-tailed Student’s t test.

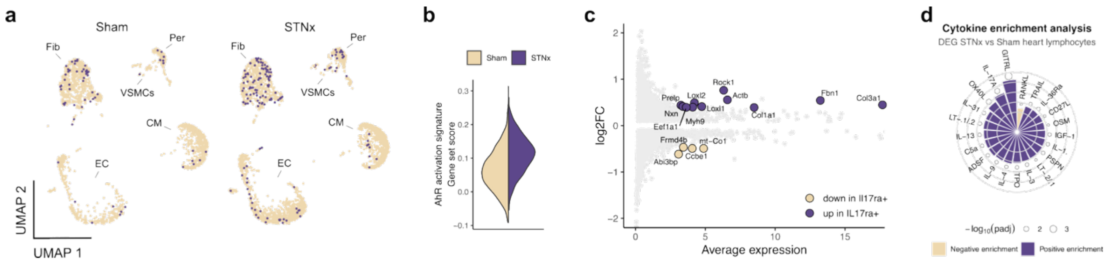

**Figure S5. Phenotype of *Il17ra* expressing cardiac fibroblasts. (a)** UMAP of previously published cardiac single nuclei (sn) RNA-sequencing data (Kaesler *et al.*) from control (Sham, left) and subtotal nephrectomized (STNx, right) mice colored according to *Il17ra* expression. **(b)** AhR activation signature in cardiac fibroblasts from Sham and STNx mice. **(c)** Differentially expressed genes in *Il17ra*-positive and -negative CKD fibroblasts. **(d)** Cytokine enrichment analysis of cardiac lymphocytes ($T_{reg}$) in STNx compared to Sham mice.

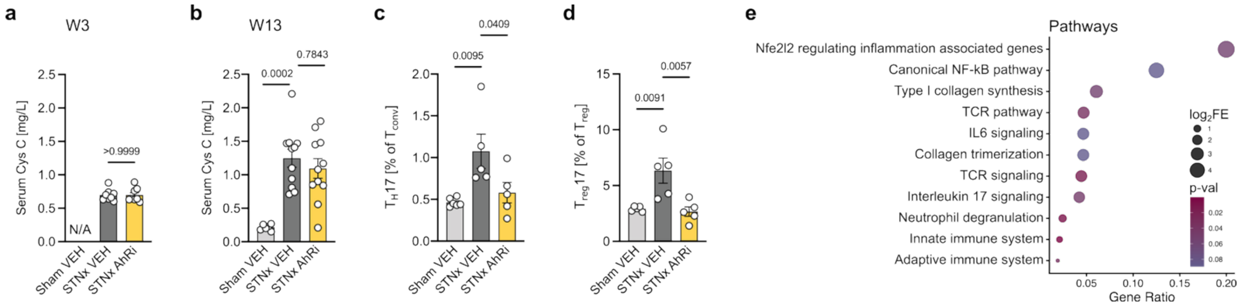

**Figure S6. Pharmacological inhibition of the AhR ameliorates kidney disease-driven inflammation.** 129/Sv mice underwent two stage subtotal nephrectomy (STNx) or sham surgery. **(a)** STNx mice were randomized two weeks after complete surgery for treatment with BAY 2416964 (AhRi) or vehicle (VEH) according to plasma cystatin C (Cys C) levels. **(b)** Cys C at study endpoint, after 9 weeks of treatment. IL-17A expression in splenic **(c)** $T_{conv}$ ($CD45^+$ $CD3^+$ $TCR\gamma\delta^-$ $CD4^+$ $CD8^-$ $FoxP3^-$) and **(d)** $T_{reg}$ ($CD45^+$ $CD3^+$ $TCR\gamma\delta^-$ $CD4^+$ $CD8^-$ $FoxP3^+$). **(e)** Significantly enriched biological pathways in the cardiac transcriptome (STNx AhRi vs. STNx). Data are shown as mean ± SEM. Each dot represents one mouse. Statistical analyses were performed using one-way ANOVA or Kruskal-Wallis test followed by Tukey's or Dunn's *post hoc* correction, as appropriate, unless otherwise indicated. Gene set statistical analyses were performed using Fisher's exact test.

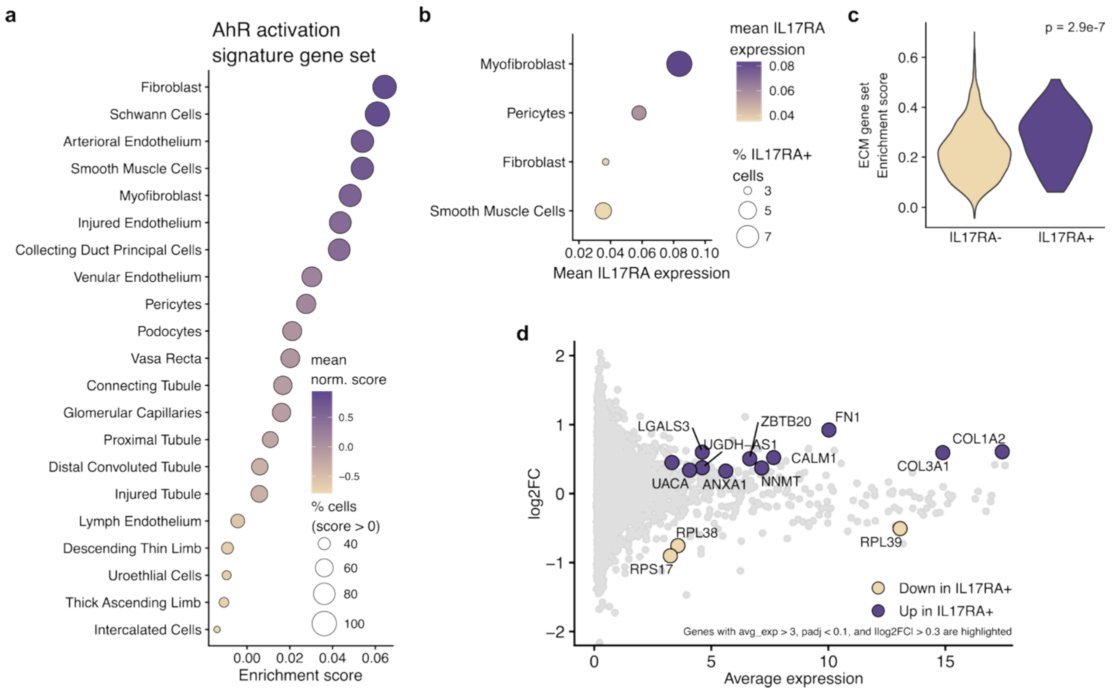

**Figure S7. Phenotype of *IL17RA* expressing human fibroblasts in kidney tissue from patients with chronic kidney disease. (a)** Analysis of previously published kidney single cell (sc) RNA-sequencing data (Kuppe *et al.*) from patients with CKD and HC were analyzed for AhR activation gene signatures. **(b)** *IL17RA* expression in kidney cell types. **(c)** Extracellular matrix (ECM) gene set expression in *IL17RA*-positive versus -negative myofibroblasts. **(d)** Differentially expressed genes (DEG) in *IL17RA*-positive versus -negative fibroblasts and myofibroblasts.

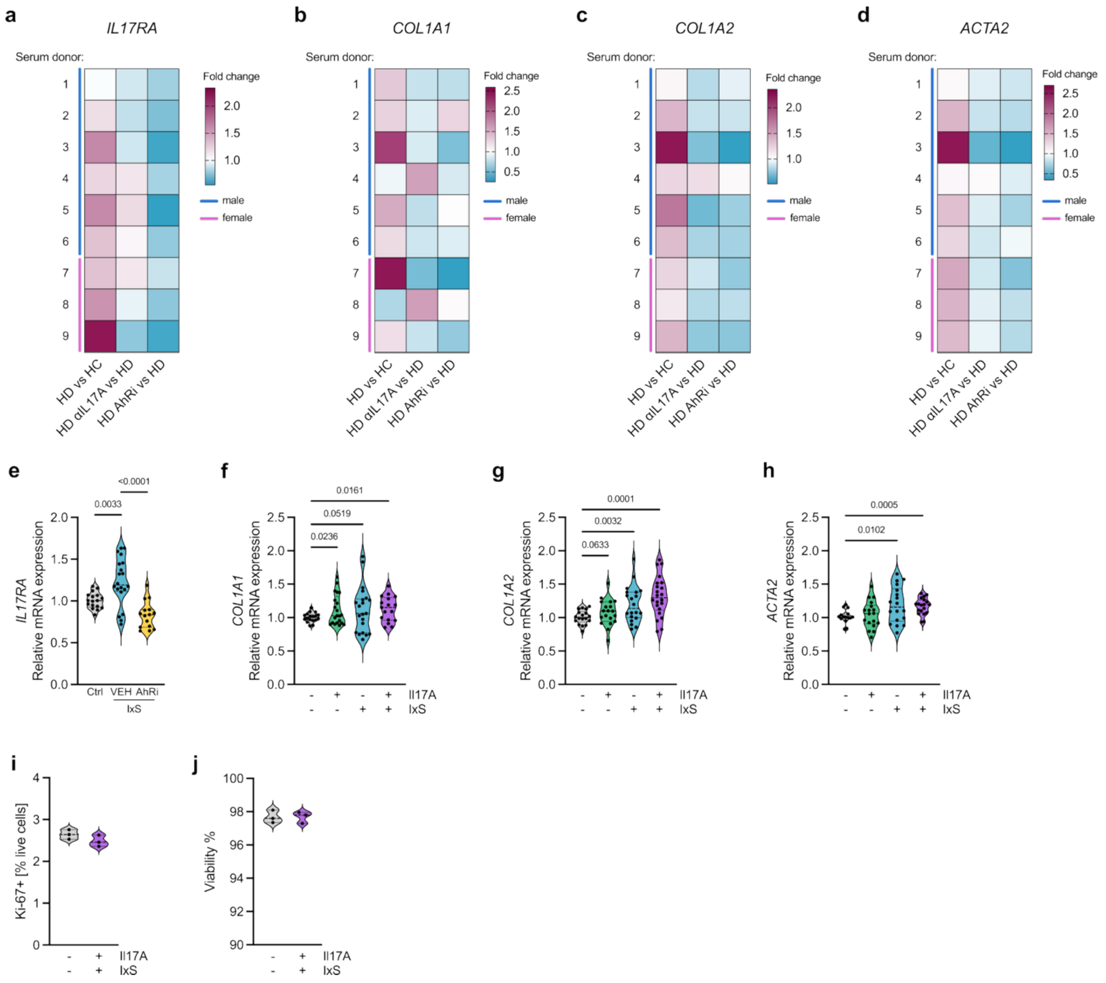

**Figure S8. Pro-fibrotic effects of indoxyl sulfate and IL-17A on human primary cardiac fibroblasts.** Serum from healthy controls (HC) and from CKD patients treated with hemodialysis (HD) was used to incubate human primary fibroblasts addition of anti-IL-17A (Secukinumab, αIL17A) or AhRi and analyzed for **(a)** *IL17RA*, **(b)** *COL1A1*, **(c)** *COL1A2*, and **(d)** *ACTA2* expression. Heatmaps show effects of individual patient serum. **(e)** Human primary cardiac fibroblasts were incubated with 100µM IxS with and without 1 µM AhRi and *IL17RA* expression was analyzed. Human primary cardiac fibroblasts were incubated with IxS (100 µM), IL-17A (1 ng/mL), or the combination of both and **(f)** *COL1A1*, **(g)** *COL1A2*, and **(h)** *ACTA2* expression were analyzed. Human primary cardiac fibroblasts were incubated with the combination of IxS (100 µM) and IL-17A (1 ng/mL) and **(i)** Ki67 expression and **(j)** viability were analyzed. **(b-e)** Primary cells from two biological replicates were treated with HC serum (n = 4 donors) or HD serum (n = 9 donors). Cells and serum were sex-matched, and for each serum donor three technical replicates were included. **(e-h)** Primary cells from two biological replicates were treated in three independent runs with the indicated substances. Within each run, three technical replicates were included. Each dot represents one well. Statistical analyses were performed using one-way ANOVA or Kruskal-Wallis test followed by Tukey's or Dunn's *post hoc* correction, as appropriate.

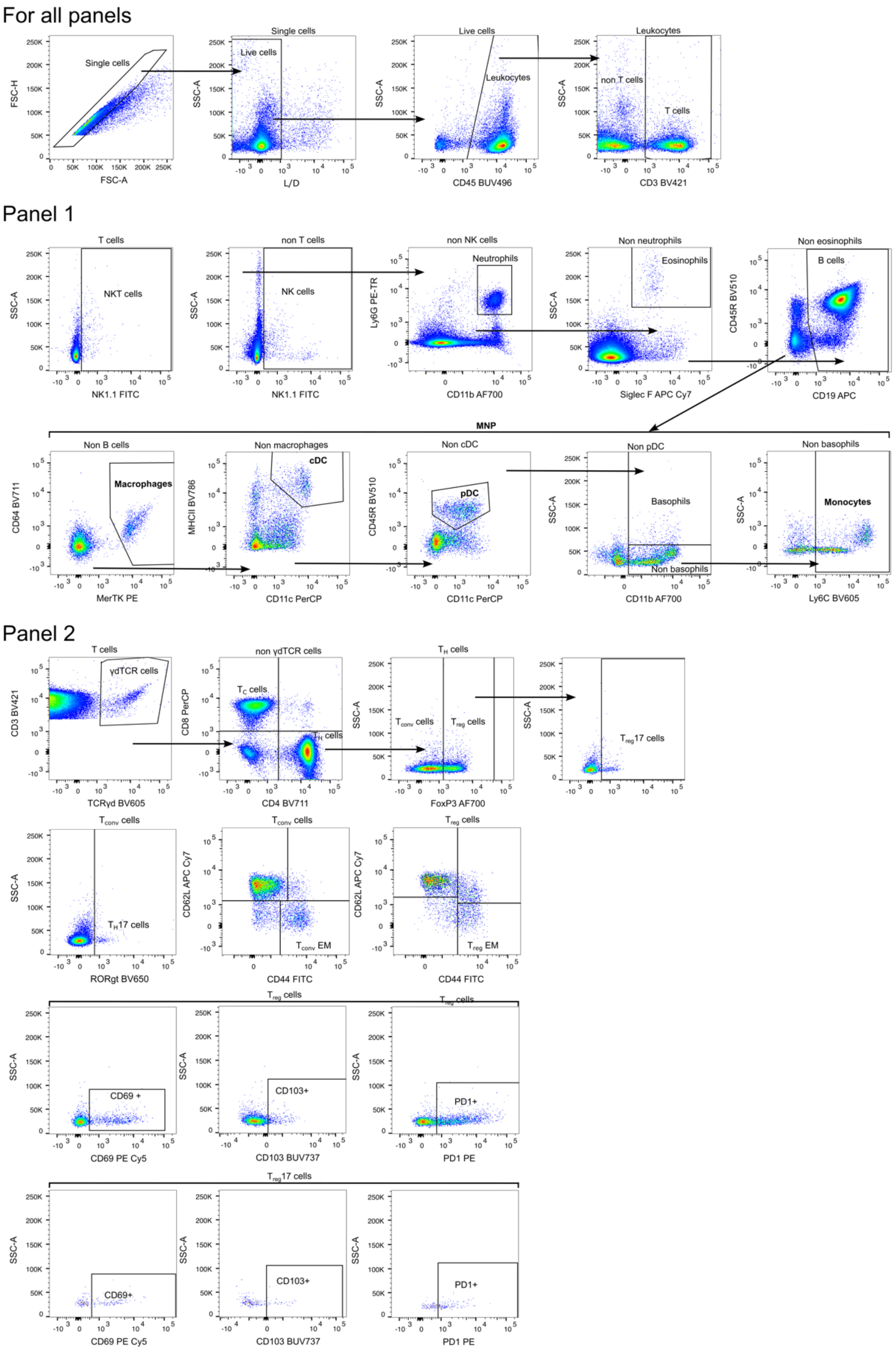

**Figure S9A. Gating strategy for mouse flow cytometry analysis of panel 1 and panel 2.**

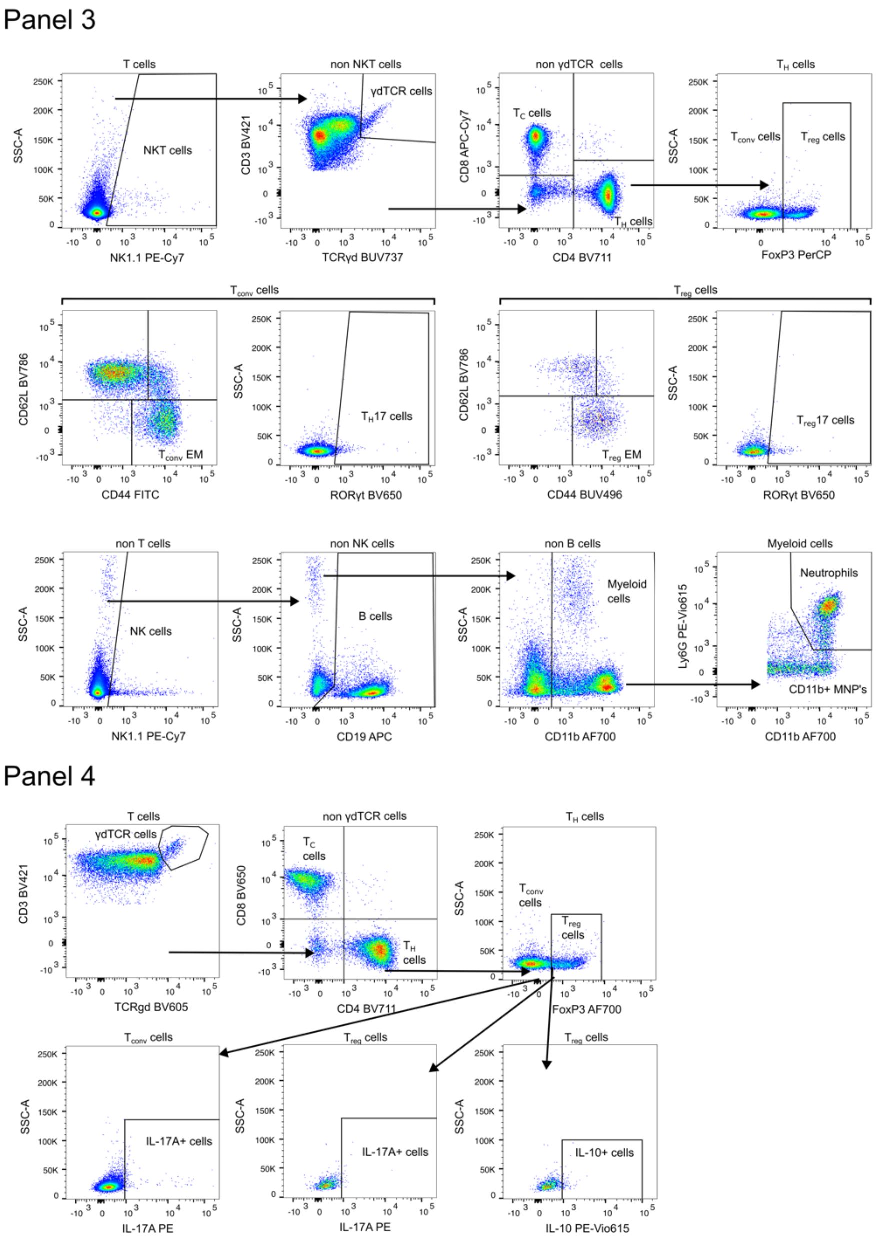

**Figure S9B. Gating strategy for mouse flow cytometry analysis of panel 3 and panel 4.**

### Supplementary tables

**Table S1: Echocardiographic parameters of the Abx trial.**

Abbreviations: LVM - left-ventricular mass, TL - tibia length, LVPWd - left-ventricular posterior wall tickness end-diastole, LVAWd - left-ventricular anterior wall thickness end-diastole, IVSd - interventricular septum thickness end-diastole, ESV - end-systolic volume, EDV - end-diastolic volume, SV - stroke volume, CO - cardiac output, EF - left-ventricular ejection fraction, FS - left-ventricular fractional shortening, IVCT - isovolumic contraction time, IVRT - isovolumic relaxation time, ET - ejection time, E - mitral E wave, A - mitral A wave, e' - mitral e' wave, GLS - left-ventricular global longitudinal strain, GRS - left-ventricular global radial strain, GCS - left-ventricular global circumferential strain.

| **Parameter** | **Sham** | **Sham Abx** | **STNx** | **STNx Abx** |
|---|---|---|---|---|
| LVM/TL (g/m) | 7.69 ± 1.03 | 6.54 ± 1.26 | 10.12 ± 1.79 | 6.55 ± 1.38 |
| LVPWd (mm) | 0.87 ± 0.08 | 0.74 ± 0.09 | 1.19 ± 0.19 | 0.89 ± 0.12 |
| LVAWd (mm) | 1.13 ± 0.14 | 1.18 ± 0.16 | 1.51 ± 0.21 | 1.34 ± 0.12 |
| IVSd (mm) | 1.00 ± 0.10 | 0.93 ± 0.10 | 1.31 ± 0.15 | 1.04 ± 0.20 |
| ESV (µL) | 35.84 ± 9.17 | 28.18 ± 9.07 | 32.05 ± 22.71 | 15.04 ± 7.61 |
| EDV (µL) | 78.07 ± 14.89 | 63.79 ± 10.68 | 67.25 ± 29.26 | 43.95 ± 10.36 |
| SV (µL) | 42.23 ± 6.36 | 35.61 ± 3.78 | 35.20 ± 10.57 | 28.91 ± 5.57 |
| CO (mL/min) | 18.27 ± 2.00 | 14.27 ± 1.83 | 16.93 ± 5.01 | 13.03 ± 3.15 |
| EF (%) | 54.53 ± 3.71 | 56.55 ± 7.23 | 62.93 ± 10.25 | 67.63 ± 12.34 |
| FS (%) | 28.08 ± 2.45 | 29.29 ± 4.69 | 34.01 ± 7.50 | 37.96 ± 11.10 |
| IVCT (ms) | 24.26 ± 4.40 | 17.45 ± 2.50 | 23.23 ± 8.35 | 17.73 ± 2.60 |
| IVRT (ms) | 18.84 ± 2.71 | 21.61 ± 2.75 | 22.09 ± 2.20 | 19.74 ± 3.13 |
| ET (ms) | 45.99 ± 3.83 | 45.64 ± 2.51 | 41.29 ± 6.60 | 41.65 ± 4.95 |
| E/A | 1.48 ± 0.27 | 1.41 ± 0.34 | 1.23 ± 0.21 | 1.25 ± 0.19 |
| E/e' | 21.58 ± 3.65 | 24.20 ± 3.65 | 33.47 ± 7.76 | 27.49 ± 6.24 |
| GLS (%) | -16.62 ± 1.63 | -17.50 ± 1.63 | -13.96 ± 2.69 | -19.41 ± 3.56 |
| GRS (%) | 22.32 ± 1.47 | 23.07 ± 1.61 | 19.45 ± 3.98 | 23.48 ± 4.60 |
| GCS (%) | -20.42 ± 2.03 | -21.86 ± 2.45 | -19.56 ± 4.58 | -23.56 ± 4.69 |

**Table S2: Echocardiographic parameters of the IxS trial.**

Abbreviations: LVM - left-ventricular mass, TL - tibia length, LVPWd - left-ventricular posterior wall tickness end-diastole, IVSd - interventricular septum thickness end-diastole, ESV - end-systolic volume, EDV - end-diastolic volume, SV - stroke volume, CO - cardiac output, EF - left-ventricular ejection fraction, FS - left-ventricular fractional shortening, IVCT - isovolumic contraction time, IVRT - isovolumic relaxation time, ET - ejection time, E - mitral E wave, A - mitral A wave, e' - mitral e' wave.

| | **HFD / L-NAME** | |
|---|---|---|
| **Parameter** | **KCl** | **IxS** |
| LVM/TL (g/m) | 6.82 ± 0.82 | 7.47 ± 1.05 |
| LVPWd (mm) | 1.04 ± 0.14 | 1.00 ± 0.10 |
| IVSd (mm) | 1.01 ± 0.13 | 1.02 ± 0.11 |
| ESV (µL) | 28.96 ± 7.61 | 27.30 ± 5.74 |
| EDV (µL) | 57.87 ± 11.18 | 58.14 ± 8.40 |
| SV (µL) | 30.53 ± 6.14 | 30.13 ± 5.84 |
| CO (mL/min) | 12.60 ± 2.94 | 13.22 ± 1.93 |
| EF (%) | 53.78 ± 10.17 | 52.69 ± 8.98 |
| FS (%) | 28.55 ± 7.54 | 27.44 ± 6.95 |
| IVCT (ms) | 17.00 ± 1.67 | 16.33 ± 1.00 |
| IVRT (ms) | 16.18 ± 1.51 | 16.33 ± 1.35 |
| ET (ms) | 47.94 ± 3.81 | 47.83 ± 4.52 |
| E/A | 2.01 ± 0.76 | 1.62 ± 0.38 |
| E/e' | 28.82 ± 6.99 | 32.52 ± 6.48 |

**Table S3: Echocardiographic parameters of the AhRi trial.**

Abbreviations: LVM - left-ventricular mass, TL - tibia length, LVPWd - left-ventricular posterior wall tickness end-diastole, LVAWd - left-ventricular anterior wall thickness end-diastole, IVSd - interventricular septum thickness end-diastole, ESV - end-systolic volume, EDV - end-diastolic volume, SV - stroke volume, CO - cardiac output, EF - left-ventricular ejection fraction, FS - left-ventricular fractional shortening, IVCT - isovolumic contraction time, IVRT - isovolumic relaxation time, ET - ejection time, E - mitral E wave, A - mitral A wave, e' - mitral e' wave, GLS - left-ventricular global longitudinal strain, GRS - left-ventricular global radial strain, GCS - left-ventricular global circumferential strain.

| Parameter | Sham VEH | STNx VEH | STNx AhRi |
|---|---|---|---|
| LVM/TL (g/m) | 8.03 ± 1.43 | 10.29 ± 2.05 | 8.45 ± 1.91 |
| LVPWd (mm) | 0.83 ± 0.17 | 1.01 ± 0.18 | 1.13 ± 0.27 |
| LVAWd (mm) | 1.23 ± 0.14 | 1.50 ± 0.27 | 1.53 ± 0.18 |
| IVSd (mm) | 1.15 ± 0.17 | 1.47 ± 0.18 | 1.51 ± 0.22 |
| ESV (µL) | 76.48 ± 10.32 | 74.30 ± 16.52 | 57.54 ± 11.76 |
| EDV (µL) | 34.84 ± 10.20 | 37.28 ± 14.78 | 27.78 ± 11.65 |
| SV (µL) | 41.64 ± 3.75 | 37.02 ± 6.70 | 29.76 ± 8.27 |
| CO (mL/min) | 18.19 ± 2.27 | 17.10 ± 3.67 | 14.13 ± 4.72 |
| EF (%) | 55.35 ± 7.64 | 51.53 ± 13.13 | 52.01 ± 8.50 |
| FS (%) | 28.76 ± 4.85 | 26.80 ± 9.52 | 26.39 ± 5.34 |
| IVCT (ms) | 18.46 ± 2.36 | 18.30 ± 4.28 | 16.44 ± 2.60 |
| IVRT (ms) | 19.61 ± 3.01 | 22.94 ± 5.63 | 23.22 ± 6.00 |
| ET (ms) | 45.79 ± 3.17 | 41.57 ± 5.28 | 40.47 ± 7.40 |
| E/A | 1.55 ± 0.25 | 1.25 ± 0.11 | 1.23 ± 0.34 |
| E/e' | 24.54 ± 3.02 | 32.35 ± 6.49 | 22.76 ± 2.71 |
| GLS (%) | -17.65 ± 2.41 | -14.95 ± 2.08 | -17.18 ± 3.38 |
| GRS (%) | 25.00 ± 6.16 | 18.27 ± 4.38 | 23.73 ± 6.95 |
| GCS (%) | -20.95 ± 3.18 | -16.43 ± 3.68 | -20.66 ± 8.36 |

**Table S4: Baseline characteristics of serum donors CKD stage 5D and healthy controls.**

| | **CKD G5D (HD)** (n = 10) | **Healthy controls (HC)** (n = 5) |
|---|---|---|
| Sex | | |
| Female | 3 | 2 |
| Male | 7 | 3 |
| Age (years) | 59.9 ± 21.0 | 36.3 ± 9.7 |
| BMI (kg/m^2) | 27.4 ± 6.5 | 22.3 ± 2.9 |
| Dialysis vintage (years) | 4.3 ± 5.2 | N/A |
| Underlying disease | | N/A |
| Diabetic nephropathy | 2 | |
| Acute tubular necrosis | 1 | |
| Autosomal recessive polycystic kidney disease | 1 | |
| Bartter syndrome | 1 | |
| Glomerulonephritis | 1 | |
| Unknown | 4 | |

**Table S5: Baseline characteristics of the UK Biobank participants stratified by eGFR.**

Data are presented as mean (standard deviation) for continuous variables and n (%) for categorical variables. Significance was assessed using linear regression for continuous and $\chi^2$ tests for categorical variables.

| | **eGFR < 60 (n=21,478)** | **eGFR > 60 (n=480,913)** | **p-value** |
|---|---|---|---|
| Cystatin C | 1.38 (0.39) | 0.88 (0.12) | <0.001 |
| Sex (male) | 12,660 (58.9%) | 216,408 (45.0%) | <0.001 |
| BMI | 30.70 (6.04) | 27.29 (4.69) | <0.001 |
| Age | 62.14 (6.11) | 56.28 (8.08) | <0.001 |
| HFpEF | 5,497 (25.6%) | 27,983 (5.8%) | <0.001 |
| All-cause death | 6,371 (29.7%) | 38,128 (7.9%) | <0.001 |
| Systolic blood pressure | 141.52 (19.41) | 137.71 (18.60) | <0.001 |
| Hip circumference | 108.54 (12.00) | 103.17 (9.04) | <0.001 |
| Whole body fat mass | 30.39 (12.02) | 24.62 (9.37) | <0.001 |
| Pulse pressure | 59.20 (15.35) | 55.45 (13.55) | <0.001 |
| Weight | 87.65 (18.74) | 77.63 (15.67) | <0.001 |
| Pulse rate | 71.46 (13.05) | 69.33 (11.17) | <0.001 |
| Diastolic blood pressure | 82.32 (10.83) | 82.25 (10.11) | 0.341 |
| Whole body fat-free mass | 57.18 (11.89) | 53.05 (11.45) | <0.001 |
| Waist circumference | 100.79 (14.67) | 89.84 (13.24) | <0.001 |
| Standing height | 168.92 (9.55) | 168.42 (9.27) | <0.001 |
| Glucose Lab | 5.43 (1.81) | 5.11 (1.21) | <0.001 |
| Alanine aminotransferase | 25.13 (16.47) | 23.47 (14.06) | <0.001 |
| Calcium - blood assay | 2.38 (0.11) | 2.38 (0.09) | 0.003 |
| Alkaline phosphatase | 96.44 (38.88) | 83.06 (25.55) | <0.001 |
| Urea | 7.15 (2.75) | 5.32 (1.24) | <0.001 |
| Glycated haemoglobin (HbA1c) | 39.61 (9.59) | 35.97 (6.58) | <0.001 |
| Cholesterol | 5.26 (1.27) | 5.71 (1.13) | <0.001 |
| HDL cholesterol | 1.23 (0.34) | 1.46 (0.38) | <0.001 |
| Red blood cell (erythrocyte) count | 4.52 (0.51) | 4.52 (0.41) | 0.120 |
| Total protein | 72.88 (4.75) | 72.49 (4.08) | <0.001 |
| Testosterone | 7.68 (5.82) | 6.50 (6.06) | <0.001 |
| Basophill percentage | 0.58 (0.70) | 0.57 (0.61) | 0.003 |
| SHBG | 46.57 (24.40) | 51.87 (27.91) | <0.001 |
| Monocyte count | 0.56 (0.34) | 0.47 (0.27) | <0.001 |
| Haemoglobin concentration | 14.13 (1.52) | 14.18 (1.23) | <0.001 |
| Basophill count | 0.04 (0.06) | 0.03 (0.05) | <0.001 |
| Lymphocyte count | 2.06 (2.04) | 1.96 (1.12) | <0.001 |
| Red blood cell (erythrocyte) distribution width | 13.98 (1.32) | 13.47 (0.96) | <0.001 |

| | | | |
|---|---|---|---|
| Haematocrit percentage | 41.11 (4.38) | 41.08 (3.51) | 0.320 |
| Neutrophill count | 4.91 (1.76) | 4.20 (1.40) | <0.001 |
| Lymphocyte percentage | 26.38 (8.58) | 29.03 (7.43) | <0.001 |
| Eosinophill count | 0.22 (0.18) | 0.17 (0.14) | <0.001 |
| Aspartate aminotransferase | 28.21 (14.07) | 26.14 (10.46) | <0.001 |
| C-reactive protein | 5.10 (7.16) | 2.48 (4.14) | <0.001 |
| White blood cell (leukocyte) count | 7.79 (2.91) | 6.84 (2.07) | <0.001 |
| Triglycerides | 2.16 (1.15) | 1.73 (1.02) | <0.001 |
| Gamma glutamyltransferase | 50.55 (64.97) | 36.76 (40.56) | <0.001 |
| Lipoprotein A | 44.37 (48.41) | 44.66 (49.25) | 0.451 |
| Monocyte percentage | 7.39 (3.63) | 7.05 (2.65) | <0.001 |
| Creatinine Lab | 99.18 (51.29) | 71.02 (14.08) | <0.001 |
| Neutrophill percentage | 62.81 (9.88) | 60.79 (8.45) | <0.001 |
| Platelet count | 249.06 (70.15) | 253.17 (59.54) | <0.001 |
| Eosinophill percentage | 2.85 (2.19) | 2.56 (1.86) | <0.001 |
| Coronary artery disease | 1,906 (8.9%) | 10,396 (2.2%) | <0.001 |
| Essential (primary) hypertension | 6,281 (29.2%) | 34,121 (7.1%) | <0.001 |
| Heart failure | 3,981 (18.5%) | 18,279 (3.8%) | <0.001 |
| Myocardial infarction | 1,307 (6.1%) | 6,505 (1.4%) | <0.001 |
| Atrial fibrillation and flutter | 1,215 (5.7%) | 7,626 (1.6%) | <0.001 |
| Type 2 diabetes | 2,495 (11.6%) | 12,800 (2.7%) | <0.001 |
| Sleep apnoea | 307 (1.4%) | 2,172 (0.5%) | <0.001 |
| Cardiomyopathy | 183 (0.9%) | 434 (0.1%) | <0.001 |
| Cancer | 2,888 (13.4%) | 35,719 (7.4%) | <0.001 |
| Betablocker | 6,039 (28.1%) | 50,132 (10.4%) | <0.001 |
| Ca-channel blocker | 5,172 (24.1%) | 42,159 (8.8%) | <0.001 |
| Cholesterol lowering medication | 8,912 (41.5%) | 77,946 (16.2%) | <0.001 |
| Vitamin C | 1,610 (7.5%) | 41,875 (8.7%) | <0.001 |
| Calcium | 1,555 (7.2%) | 33,304 (6.9%) | 0.083 |
| Thiazide diuretics | 4,770 (22.2%) | 38,741 (8.1%) | <0.001 |
| ACE-inhibitor | 7,537 (35.1%) | 56,149 (11.7%) | <0.001 |
| ARB | 3,229 (15.0%) | 20,097 (4.2%) | <0.001 |
| Loop diuretics | 2,352 (11.0%) | 7,331 (1.5%) | <0.001 |
| Metformine | 893 (4.2%) | 5,176 (1.1%) | <0.001 |
| Aspirin | 7,464 (34.8%) | 70,193 (14.6%) | <0.001 |
| Insulin | 1,056 (4.9%) | 4,554 (0.9%) | <0.001 |

**Table S6: Overview of primers and probes used in this study.**

Abbreviations: Pro - prokaryotes, Eu - eukaryotes, F - forward primer, R - reverse primer, P - probe, IDT - Integrated DNA Technologies, Thermo - Thermo Fisher Scientific.

| Gene | Species | Type | Sequence | nM | Vendor |
|---|---|---|---|---|---|
| *16S* | Pro | Forward | ACTCCTACGGGAGGCAGCAGT | 450 | IDT |
| | | Reverse | GTATTACCGCGGCTGCTGGCAC | 450 | IDT |
| *18S* | Eu | Forward | CTGAGAAACGGCTACCACATC | 75 | IDT |
| | | Reverse | GCCTCGAAAGAGTCCTGTATTG | 75 | IDT |
| | | Probe | VIC-AAATTACCCACTCCCGACCCGG-QSY | 100 | Thermo |
| *Abi3bp* | mouse | Forward | TTCACCAGTCCTGAAGTGCGAC | 300 | IDT |
| | | Reverse | CAGGCTTGACTCTTGGCGGTTT | 300 | IDT |
| *ACTA2* | human | Forward | CTATGCCTCTGGACGCACAACT | 300 | IDT |
| | | Reverse | CAGATCCAGACGCATGATGGCA | 300 | IDT |
| *Acta2* | mouse | Forward | GACTCTCTTCCAGCCATCTTTC | 300 | IDT |
| | | Reverse | GACAGGACGTTGTTAGCATAGA | 300 | IDT |
| | | Probe | JUN-CGGGCATCCACGAAACCACCTATAA -QSY | 150 | Thermo |
| *COL1A1* | human | Forward | GATTCCCTGGACCTAAAGGTGC | 300 | IDT |
| | | Reverse | AGCCTCTCCATCTTTGCCAGCA | 300 | IDT |
| *Col1a1* | mouse | Forward | CCTCAGGGTATTGCTGGACAAC | 300 | IDT |
| | | Reverse | CAGAAGGACCTTGTTTGCCAGG | 300 | IDT |
| *COL1A2* | human | Forward | CCTGGTGCTAAAGGAGAAAGAGG | 300 | IDT |
| | | Reverse | ATCACCACGACTTCCAGCAGGA | 300 | IDT |
| *Col1a2* | mouse | Forward | GTGAGTGTAGACCATCCCAATG | 450 | IDT |
| | | Reverse | TCTTCCAGCACCGTTCATATTC | 450 | IDT |
| | | Probe | FAM-AGTTGACCAAGGCTGACACGAACT-QSY | 200 | Thermo |
| *Cyp1a1* | mouse | Forward | TCCTACCTATGGGCTGACTT | 450 | IDT |
| | | Reverse | GGACAGTCTAAGCCTGAAGATG | 450 | IDT |
| | | Probe | FAM-TTCCAAGTGCAGATGCGGTCTTCT-QSY | 200 | Thermo |
| *Gapdh* | mouse | Forward | TCTCCCTCACAATTTCCATCC | 300 | IDT |
| | | Reverse | GGGTGCAGCGAACTTTATTG | 300 | IDT |
| | | Probe | VIC-AGCCCTCCCTACTCTCTTGAATACCA-QSY | 150 | Thermo |
| *HPRT1* | human | Forward | CATTATGCTGAGGATTTGGAAAGG | 300 | IDT |
| | | Reverse | CTTGAGCACACAGAGGGCTACA | 300 | IDT |
| *Hprt* | mouse | Forward | CTGGTGAAAAGGACCTCTCGAAG | 300 | IDT |
| | | Reverse | CCAGTTTCACTAATGACACAAACG | 300 | IDT |
| *IL17RA* | human | Forward | TCATCGTCTGCATGACCTGGAG | 300 | IDT |
| | | Reverse | GGCTGAGTAGATGATCCAGACC | 300 | IDT |
| *Il17ra* | mouse | Forward | CTGTATGACCTGGAGGCTTTCTG | 300 | IDT |
| | | Reverse | CGAGTAGACGATCCAGACCTTC | 300 | IDT |

**Table S7: Overview of mouse antibodies used in this study.**

Panels were stained in the following trials and organs: 1 - Abx trial; spleen. 2 - Abx trial; spleen, PBC, intestine. 3 - Abx trial; heart and IxS and AhRi trials; spleen. 4 - Abx, IxS, AhRi trials; spleen. 5 - Murine $T_H$17 assay. 6 - Murine cardiac fibroblasts.

| Target | Fluorophore | Clone | Manufacturer | Reference | Dil. | Panel |
|---|---|---|---|---|---|---|
| CD3 | BV421 | 17A2 | BD Biosciences | 564008 | 1:50 | 1,2,3,4 |
| CD4 | BV711 | RM4-5 | BioLegend | 100549 | 1:100 | 2,3,4 |
| CD4 | VioGreen | GK1.5 | Miltenyi Biotec | 130-123-899 | 1:100 | 5 |
| CD8a | APC-Vio770 | 53-6.7 | Miltenyi Biotec | 130-123-280 | 1:400 | 3 |
| CD8a | BV650 | 53-6.7 | BioLegend | 100741 | 1:100 | 4 |
| CD8a | PerCP-Cy5.5 | 53-6.7 | invitrogen | 45-0081-82 | 1:100 | 2 |
| CD11b | AF700 | M1/70 | BD Biosciences | 557960 | 1:100 | 1,3 |
| CD11c | PerCP-Cy5.5 | N418 | BioLegend | 117328 | 1:50 | 1 |
| CD19 | APC | REA749 | Miltenyi Biotec | 130-111-884 | 1:200 | 3 |
| CD25 | PE | REA568 | Miltenyi Biotec | 130-120-696 | 1:200 | 5 |
| CD31 | APC | REA784 | Miltenyi Biotec | 130-111-541 | 1:100 | 6 |
| CD44 | BB770 | IM7 | Miltenyi Biotec | 566506 | 1:200 | 5 |
| CD44 | FITC | IM7 | BD Biosciences | 553133 | 1:100 | 2,3 |
| CD45 | BUV495 | 30-F11 | BD Biosciences | 749889 | 1:400 | 1,2,3,4,6 |
| CD45R | BV510 | RA3-6B2 | BioLegend | 103248 | 1:200 | 1 |
| CD62L | APC-Cy7 | MEL-14 | BioLegend | 104428 | 1:200 | 2 |
| CD62L | BV421 | MEL-14 | BD Biosciences | 562910 | 1:200 | 5 |
| CD62L | BV786 | MEL-14 | BioLegend | 104440 | 1:200 | 3 |
| CD64 | BV711 | X54-5/7.1 | BioLegend | 139311 | 1:50 | 1 |
| CD69 | PE-Cy5 | H1.2F3 | BioLegend | 104510 | 1:100 | 2 |
| CD103 | BUV737 | 2E7 | BD Biosciences | 749393 | 1:200 | 2 |
| CD140a | PE | APA5 | Miltenyi Biotec | 130-102-502 | 1:20 | 6 |
| Foxp3 | AF700 | FJK-16s | invitrogen | 56-5773-82 | 1:100 | 2,4 |
| Foxp3 | PerCP-Cy5.5 | FJK-16s | invitrogen | 45-5773-82 | 1:100 | 3 |
| IL10 | PE-Dazzle594 | JES5-16E3 | BioLegend | 505034 | 1:100 | 4 |
| IL17A | PE | eBio17B7 | invitrogen | 12-7177-81 | 1:100 | 4,5 |
| Ly6C | BV605 | AL-21 | BD Biosciences | 563011 | 1:100 | 3 |
| Ly6G | PE-Vio615 | REA526 | Miltenyi Biotec | 130-123-029 | 1:200 | 1,3 |
| MHCII | BV786 | M5/114.15.2 | BioLegend | 107645 | 1:400 | 1 |
| NK1.1 | PE-Vio770 | REA1162 | Miltenyi Biotec | 130-120-509 | 1:100 | 3 |
| NK1.1 | VB-B515 | REA1162 | Miltenyi Biotec | 130-120-503 | 1:50 | 1 |
| PD1 | PE | 29F.1A12 | BioLegend | 1345206 | 1:400 | 2 |
| RORγt | BV650 | Q31-378 | BD Biosciences | 564722 | 1:50 | 2,3 |
| Siglec F | APC-Vio770 | REA798 | Miltenyi Biotec | 130-112-177 | 1:200 | 1 |
| TCRγδ | BUV737 | GL3 | BD Biosciences | 748991 | 1:100 | 3 |
| TCRγδ | BV605 | GL3 | BioLegend | 118129 | 1:50 | 2,4 |

**Table S8: Overview of human antibodies used in this study.**

| Target | Fluorophore | Clone | Manufacturer | Reference | Dil. |
|---|---|---|---|---|---|
| *Human $T_H$17 assay:* | | | | | |
| CD4 | VioGreen | VIT4 | Miltenyi Biotec | 130-113-221 | 1:50 |
| CD25 | PE | BC96 | BioLegend | 302606 | 1:200 |
| CD45RO | FITC | UCHL1 | BioLegend | 304242 | 1:400 |
| CD127 | BV421 | A019D5 | BioLegend | 351310 | 1:200 |
| IL17A | APC-Vio770 | REA1063 | Miltenyi Biotec | 130-118-249 | 1:50 |
| *Human cardiac fibroblasts:* | | | | | |
| Ki-67 | PE Vio-770 | REA183 | Miltenyi Biotec | 130-120-419 | 1:100 |

### Supplementary methods

#### Animal studies

All experiments were performed in accordance with the German/European law for animal protection. Experimental protocols were approved by the local ethic authorities under the G0190/21 and G0019/21 licenses. For all *in vivo* experiments, male 129/Sv mice (wild type [WT]; strain 129S2/SvPasOrlRj) were obtained from Janvier Labs, France, and housed at 12-hour day:night cycle with free access to water and food (Chow diet, Ssniff, V1124-300) in the Preclinical Research Center (PRC) of the Max-Delbrück-Center for Molecular Medicine (MDC). Mice were habituated for 2 weeks before use. Individual mice served as experimental units, and all animals within a cage received the same treatment.

For the Abx trial, mice were distributed according to body weight (BW) to four groups prior to treatment (STNx ± Abx and Sham ± Abx). The Abx experiment consisted of two independent cohorts. A longitudinal cohort was designed to evaluate dynamics in kidney function, metabolomics and immunity in STNx groups (n=11 per group) and Shams (n=7 per group) across timepoints (w1, w3, w8, w13). Each timepoint had its own independent cohort to collect biomaterial. A second cohort was designed for terminal assessments, focused on organ damage and tissue level responses in STNx groups (n=12) and Shams (n=7). Both cohorts were exploratory, thereby no prior sample size calculation was performed. Endpoint analyzes were performed combining both cohorts if applicable. Animals were excluded in cases of surgical failure or failure to reach minimum kidney function thresholds. Animals were sacrificed during the course of the study if BW dropped under 20% or they showed clear signs of pain or discomfort. The longitudinal cohort resulted in: w1 - Sham n=7; Abx n=6; STNx n= 6; STNx Abx n= 6. w3 - Sham n=6; Abx n= 7; STNx n=10; STNx Abx n=7. w8 - Sham n=5; Abx n=5; STNx n=6; STNx Abx n=6. w13 - Sham n=6; Abx n=4; STNx n=6; STNx Abx n=6. The endpoint cohort resulted in: Sham n=7; Abx n=7; STNx n=11; STNx Abx n=10. The telemetry cohort resulted in STNx n=7 STNx, STNx Abx n=7).

For the HFD/L-NAME trial, mice were assigned according to BW and cage to receive either potassium indoxyl sulfate (IxS) exposure (n=16) or control (KCl; n=16) over the drinking water *ad libitum.* Sample size was calculated using G Power according to a partial effect size of the alleviation of the STNx-induced diastolic dysfunction by Abx. During the experiments n=3 from the control group were taken out of the experiment due to prespecified criteria.

For the AhRi trial, mice underwent STNx (n=24) or Sham (n=6) surgery. Sample size was calculated using G Power according to the effect size of the alleviation of the STNx-induced cardiac fibrosis by Abx. The same exclusion criteria were followed as in the Abx trial. To ensure equal kidney dysfunction before treatment, mice were equally distributed into 2 groups according to BW and plasma CysC two weeks after completed STNx surgery, and assigned to receive either AhRi (n=10) or vehicle (n=9). During the experiments n=4 vehicle and n=2 AhRi-treated animals were taken out of the experiment due to prespecified criteria and analyzed at an earlier time point.

Across all experiments, blinding during *in vivo* procedures was not possible due to model-specific features; however, all outcome measurements and data analysis were performed blindly.

<u>Subtotal nephrectomy</u>

Mice between 10 and 12 weeks old were anaesthetized with isoflurane (2.5% isoflurane-air) and subjected to subtotal nephrectomy operation (STNx). Briefly, the left kidney was exposed, and the lower and upper poles were excised using a cauterizer (Surtron 160, LED SpA). The adrenal gland was separated from the kidney to preserve its function. 2 weeks later, a total nephrectomy of the right kidney was performed. Sham-operated mice underwent a simple flank incision and sutures. All animals received pre-emptive buprenorphine (30 mg/kg, intraperitoneal (i.p.) injection) 30 min before surgery. Post surgery, analgesia consisted of metamizole (800 mg/kg/24h) administered over the drinking water for 3 days.

Antibiotic treatment

Mice were given an antibiotic cocktail (all obtained individually from Thermofisher) containing ampicillin (0.5 g/L), vancomycin hydrochloride (0.5 g/L), neomycin sulfate (1 g/L) and metronidazole (1 g/L) in the drinking water throughout the entire study period. 1% glucose was added into the mixture for organoleptic purposes. Control animals received only glucose in the drinking water. Treatment was started one week prior to the first operation and antibiotics were refreshed every 3 days.

AhR inhibitor treatment

Mice were stratified and randomized as described above to receive either small-molecule AhR inhibitor BAY2416964 (GlpBio) or vehicle, starting two weeks after completed STNx surgery. AhRi was formulated in a mixture of water, Solutol® HS 15 (polyethylene glycol 12-hydroxystearate; Merck) and ethanol at a 50:40:10 ratio (v/v)[1]. The vehicle control consisted of the solvent mixture alone. A daily AhRi dose of 30 mg/kg was selected based on a previously reported safe and effective regimen[1].

HFpEF model

12-week-old mice were switched to a high fat diet (HFD; 60% kcal fat, Research diets, D19032801-1.5V) and administered Nω-nitro-L-arginine methyl ester hydrochloride (L-NAME; 1 g/L, Merck) in the drinking water for 15 weeks. Mice received either potassium indoxyl sulfate (IxS; 1 g/L, Merck) or equimolar potassium chloride (KCl; 0.3 g/L, Merck) in the drinking water throughout the study. This model has been published previously[2].

**Telemetry**

In a subset of mice (n=7 STNx, n=7 STNx Abx) subcutaneous radiotelemetry transmitters (PA-C10 BP, DSI data science international, St. Paul, MN, USA) were implanted one week prior to the administration of antibiotics. Due to the length of the study, we measured blood pressure on a weekly 3-day period (except w2, which had a 2-day measurement). Means and SD per each 3-day measurement period were calculated and values with an absolute SD > 5 were excluded from downstream analysis. Longitudinal curves were generated by a non-parametric, locally weighted scatterplot smoothing (LOESS) regression. For statistical analysis, systolic and diastolic blood pressure AUCs were computed and divided by the total elapsed recording time (in minutes) to yield average blood pressure per minute. Group differences were assessed using a Wilcoxon test. Of the 14 mice initially implanted with radiotelemetry transmitters, 5 mice died perioperatively and were excluded. An additional 3 mice were removed prior to analysis due to telemetry signal instability and inconsistent blood pressure recordings. Thus, only animals that survived surgery and maintained stable telemetry signals throughout the study (n=3 per group) were included in the final analysis.

**Patient serum collection**

Patients with CKD stage 5 dependent on hemodialysis were recruited Department of Nephrology and Medical Intensive Care at Charité-Universitätsmedizin Berlin. Healthy volunteers were recruited at the

Experimental and Clinical Research Center (ECRC) of Charité-Universitätsmedizin Berlin. Ethical approval was obtained from the institutional review board of Charité–Universitätsmedizin Berlin (EA2/162/17) and all participants provided written informed consent before biomaterial was collected. Blood was obtained from patients at the start of hemodialysis. Serum was collected by centrifugation at 1500 g for 15 min at RT, and stored at -80 °C until further use. Demographic and clinical characteristics of the study cohort are summarized in Table S4.

**UK Biobank**

<u>Study participants</u>

The UK Biobank is a population-based cohort of 502,366 participants aged 40-69 years recruited between 2006 and 2010 across 22 assessment centers in England, Scotland, and Wales. Ethical approval was granted by the North West Multi-Centre Research Ethics Committee (reference 11/NW/0382), and all participants provided written informed consent. Data available as of 16 November 2023 were used for this study.

Participants with impaired kidney function were defined as those with an estimated glomerular filtration rate (eGFR) below 60 mL/min/1.73 m², corresponding to KDIGO categories G3-G5. Demographic and clinical characteristics of the study cohort are summarized in Table S5.

Identification of individuals with heart failure with preserved ejection fraction (HFpEF) followed a previously published multi-stage algorithm[3]. All HFpEF cases were required to exhibit at least one clinical symptom of heart failure in accordance with European Society of Cardiology (ESC) guidelines, supplemented by additional criteria depending on data availability. When cardiac magnetic resonance (CMR) imaging was available, participants with a left ventricular ejection fraction above 49% were considered to have preserved systolic function. When NT-proBNP measurements were available, patients exceeding the 90th percentile were classified as HFpEF, whereas those with lower values or evidence of reduced ejection fraction were excluded. For participants lacking imaging or proteomic data, HFpEF was assigned when the pre-test probability exceeded 90%.

<u>Data modalities</u>

All variables used in this study were assessed prior to or at the baseline visit. eGFR was calculated from plasma cystatin C concentrations using the equation eGFR = 74.835 / (cystatin $C^{1.333}$). Albuminuria was quantified as the urine albumin-to-creatinine ratio. Essential hypertension was defined by the presence of an ICD-10 I10 diagnosis code recorded before baseline. Proteomics profiling was available for approximately 10% of the UK Biobank participants and was performed using Olink proximity extension assays, with protein concentrations reported as $log_2$-normalized protein expression (NPX) values. For the present analysis, we focused on NT-proBNP and IL-17A, which were available for 51,578 and 50,573 participants, respectively. Smoking exposure was quantified using the UK Biobank variable "pack years of smoking" (Field-ID 20161), calculated from baseline questionnaire responses using the formula: (cigarettes per day / 20) * (age stopped smoking - age started smoking). This measure reflects cumulative smoking intensity, with higher values reflecting greater lifetime exposure.

<u>Statistical Analysis</u>

Time-to-event outcomes were evaluated using Kaplan-Meier estimators implemented in the Python package lifelines (version 0.30.0). Participants were stratified into three IL-17A groups according to the plasma IL-17A concentrations: below the 25th percentile, between the 25th and 75th percentiles, and above the 75th percentile. Survival time was defined as the interval from baseline to the occurrence of the event of interest, with censoring at the end of follow-up. Differences across IL-17A strata were assessed using the log-rank test.

To evaluate the association between IL-17A and clinical outcomes, Cox proportional hazards models were fitted. All covariates were standardized before model fitting to facilitate direct comparison of effect sizes. Cox proportional hazards models were estimated using an unpenalized framework with robust variance estimation. To ensure numerical stability, a limited grid of ridge penalisation terms ($\lambda = 0$ to 0.01) was evaluated, and the model yielding stable convergence without materially influencing effect estimates was selected. Hazard ratios (HRs) and 95% confidence intervals (CIs) were derived from the final model and displayed using forest plots.

Normality of continuous variables was assessed using the Shapiro-Wilk test. When both groups satisfied the normality assumption ($p > 0.05$), differences were evaluated using Welch's *t* test; otherwise, comparisons were performed using the non-parametric Mann-Whitney U test. For analyzes involving multiple statistical tests, p-values were adjusted using the Benjamini-Hochberg false discovery rate (FDR) procedure. All statistical analyzes were performed in Python 3.11.

**Microbiome**

Cecal DNA isolation

At different timepoints (w1, w3, w8, w13), cecal contents were collected at the time of euthanasia. Samples were immediately fresh-frozen on dry ice and kept at -80 °C. DNA was isolated from cecal samples using the Quick-DNA Miniprep Kit (Zymo Research) according to manufacturer's protocol. Absolute bacterial load was determined by qPCR for the *16S* gene using SYBR Green as described below.

Microbiome analysis

Samples were stored at -20°C and sent for 16S rRNA gene amplicon sequencing (Novogene, Cambridge, UK). The V3-V4 hypervariable region of the 16S rRNA gene was amplified using universal primers 341F (5′-CCTAYGGGRBGCASCAG-3′) and 806R (5′-GGACTACNNGGGTATCTAAT-3′), which were appended with sample-specific barcodes and Illumina sequencing adapters. Amplicons were purified using AMPure XP magnetic beads and quantified for library preparation.

Libraries were constructed using the NEBNext® Ultra™ II DNA Library Prep Kit and sequenced on an Illumina NovaSeq 6000 platform (paired-end 2×250 bp), targeting approximately 30,000 raw reads per sample. Annotation was performed in R (v4.0.3). Demultiplexed reads were processed through DADA2 pipeline (v1.25.2) and taxonomic assignment was based on the rRNA database SILVA (v138.1) with a 80% boodstrapped confidence. Only ASVs that were present in at least 20% of the samples were considered for further downstream analysis.

**Nuclear magnetic resonance (NMR) spectroscopy**

Sample preparation

Plasma samples were prepared using 3 mm SampleJet NMR tubes. 100 µL of plasma was mixed thoroughly with 78 µL of 0.1 mol/L phosphate buffer (pH 7.4) and 19.5 µL of a 0.75% (w) solution of 3-trimethylsilyl-2,2,3,3-tetradeuteropropionate (TSP; Sigma-Aldrich, Taufkirchen, Germany) in deuterium oxide, serving as an internal standard for NMR measurements. As an additional internal standard 2.5 µL of formic acid (240 mM) was added. Prepared samples were measured as described below.

Data acquisition

All NMR experiments were performed on a 600 MHz Bruker Avance III HD spectrometer, using a triple resonance ($^{1}H$, $^{13}C$, $^{15}N$, $^{2}H$ lock) helium cooled cryoprobe with z-gradient. Samples were handled by an automatic Bruker SampleJet sample changer (Bruker Biospin GmbH, Ettlingen, Germany). Tuning and matching of the probe as well as locking and shimming of the sample were performed automatically. For each sample one dimensional $^{1}H$ NMR spectra were obtained at 310 K using Carr-Purcell-Meiboom-Gill (CPMG) pulse sequence. For each spectrum, 512 scans were collected into 72k datapoints over a spectral width of 12019 Hz. An acquisition time and $T_1$ relaxation delay of 3.07 and 4 seconds per scan were used, respectively. All NMR spectra were processed with TopSpin 4.5.0. Free induction decays (FIDs) were Fourier transformed, phase corrected and subjected to exponential line broadening function of 0.3 Hz. All the chemical shifts were manually referenced to formic acid at 8.46 ppm.

To compensate for chemical shifts perturbations bucket tables were generated from the processed 1D CPMG spectra using AMIX software (version 3.9.13, Bruker, Ettlingen, Germany). The spectra over a range from $\delta$0.11 to $\delta$ 9.5 ppm were divided into the buckets at an equal width of 0.01 ppm with the regions at $\delta$4.59 - $\delta$4.77 ppm being excluded as the water peak. The spectra were not normalized.

Bucket analysis

Only buckets within the aromatic region (6.75-8.00 ppm) were retained for subsequent analyzes. For each treatment background, sham animals were used to calculate the mean intensity for every aromatic bucket, providing treatment-matched reference values. Each STNx (± Abx) sample was then compared with its corresponding sham reference by determining, for each bucket, whether its intensity exceeded the sham mean. The proportion and number of aromatic buckets classified as increased were calculated per sample and statistically compared between groups using Mann Whitney U test.

### Liquid chromatography-mass spectrometry

Tryptophan panel

Tryptophan and its downstream metabolites were quantified in serum and plasma samples. If not stated otherwise, the sample preparation was applied as stated previously[4]. Briefly, 10 µL of an aqueous stable-isotope labelled internal standard mixture was added to 50 µL of serum/ plasma. For protein precipitation samples were diluted in a ratio of 1:5 with 100% cold methanol, vortexed, and stored at -80 °C for 24 h. Samples were centrifugated at 10,000×g at 4 °C for 5 min. The supernatant was collected. The protein pellet was washed twice by addition of 200 µL of cold 80% MeOH and subsequent centrifugation at 10,000×g (washing step 1) and 12,000×g (washing step 2) at 4 °C for 5 min. For each sample, all supernatants were combined and dried using a vacuum evaporator (CombiDancer, Hettich AG, Bäch, Switzerland). All samples were reconstituted in 100 µL in water.

An ExionLC-30AD HPLC system (AB Sciex, Germany, Darmstadt) was used for the chromatographic separation of metabolites on an ACQUITY Premier HSS T3, 1.8 µm, 2.1 × 150 mm reversed-phase column (Waters, Eschborn, Germany) reversed-phase column. Gradient elution was performed with mobile phase A consisting of 0.1% formic acid in water and mobile phase B of 0.1% formic acid in acetonitrile. Metabolite detection was performed with a TripleQuad6500$^+$ (AB Sciex, Germany, Darmstadt) performing positive/ negative-switching for the ionization. Peak integration and data evaluation were carried out using SciexOS-MQ Software (Version 2.1.6, AB Sciex, Germany, Darmstadt). Results were normalized to sample volumes.

3-Indoxyl sulfate quantification

For the HFD/L-NAME trial, frozen plasma samples were thawed at RT and then 70 µL of each sample was mixed with 210 µL of a cold methanolic solution of the internal standard $d_5$-IxS (Toronto Research Chemicals, Toronto, Canada), concentrated to 1 µmol/L . Proteins were precipitated by vortexing for 1 min and subsequent storage for 60 min at -20°C. Supernatants were obtained by centrifugation at 9,400 g at 4°C for 10 min and subjected to LC-MS/MS IxS quantification applying the multiple reaction monitoring (MRM) method. Chromatographic separation was achieved on a 1290 Infinity II HPLC (Agilent Technologies, Waldbronn, Germany) equipped with a Poroshell 120 EC-C18 column (3.0 x 150 mm, 2.7 µm; Agilent Technologies) guarded by a pre-column (3.0 x 5 mm, 2.7 µm) of identical material. Water (eluent A) and acetonitrile (eluent B), both acidified with 0.1% formic acid, were pumped with 0.4 mL/min. Elution of IxS and its internal standard was achieved with a 10-min linear gradient from 5% to 90% eluent B. Total run-time was 16 min including re-equilibration of the LC system. MS/MS analyzes were carried out using an Ultivo (6465B) triple-quadrupole mass spectrometer (Agilent Technologies) operating in the negative electrospray ionization mode (ESI-). The following ion source parameters were set: sheath gas temperature, 400°C; sheath gas flow, 12 l/min of nitrogen; nebulizer pressure, 50 psi; drying gas temperature, 100°C; drying gas flow, 9 l/min of nitrogen; capillary voltage, 2.0 kV; nozzle voltage, 0.4 kV.

The following mass transitions were recorded (fragmentor voltage [FV] and collision energies [CE] in parentheses): IxS: *m/z* 212.0 → 79.8 (FV: 102 V, CE: 28 eV, quantifier), *m/z* 212.0 → 80.9 (FV: 102 V, CE: 16 eV), *m/z* 212.0 → 131.9 (FV: 102 V, CE: 20 eV), $d_5$-IxS: *m/z* 217.0 → 79.8 (FV: 82 V, CE: 28 eV, quantifier), *m/z* 217.0 → 80.9 (FV: 82 V, CE: 16 eV), *m/z* 217.0 → 137.0 (FV: 82 V, CE: 20 eV). Peak areas were determined with MassHunter Quantitative Analysis software (version 10.1, Agilent Technologies) and IxS was directly quantified via its internal standard $d_5$-IxS that was concentrated to 750 nmol/L in the samples.

**Transdermal measurement of glomerular filtration rate**

FITC-sinistrin was reconstituted with PBS (40 mg/mL) and stored at -20 °C. Mice were anaesthetized with isoflurane and prone-placed on a heat pad. After shaving, the GFR monitor was attached on the right side of the back below the ribs. The measurement was recorded for 2-3 minutes prior to intravenous FITC-sinistrin injection (60 mg/kg) via either the tail vein or retro-orbital injection to set a steady baseline. Mice were then allowed to recover from anesthesia for the measurement of FITC decay (1-2 hours). Transcutaneous GFR was calculated from FITC-sinistrin half-time using a 3-compartment model (MBLab2, Medibeacon).

**Clinical chemistry**

Clinical parameters (plasma CRP, Cystatin C) were analyzed using an AU480 clinical chemistry analyzer (Beckman Coulter).

**Echocardiography**

Echocardiography was performed according to a standard operation protocol[5]. Briefly, mice were examined on a Vevo 3100 Imaging System equipped with a 30 MHz linear transducer (MX400; FUJIFILM VisualSonics Inc., Canada). Anaesthesia was induced by 3% isoflurane (in 80% oxygen). For image acquisition, isoflurane concentration was reduced to 1-1.5%, and adjusted to maintain comparable heart rates. All images were analyzed using Vevo LAB analysis software (FUJIFILM VisualSonics Inc.). Systolic function and cardiac dimensions were assessed using B-mode and M-mode echocardiography obtained from the parasternal long-axis view and the parasternal short-axis view at the mid-papillary level. Diastolic function was evaluated using conventional pulsed-wave Doppler to measure transmitral flow in the apical four-chamber view. Tissue Doppler recordings from the septal mitral annulus were used to assess myocardial relaxation velocity. The following parameters were analyzed: E/A (early diastolic filling velocity to atrial contraction), E/e′ (early filling velocity to early diastolic annular motion), and IVRT (isovolumetric relaxation time). Strain and M-mode measurements were derived from three images per parameter, with three cardiac cycles per image, and subsequently averaged. Diastolic Doppler measurements were obtained from three independent images, each with three repeated measurements. Speckle-tracking analysis was performed using parasternal long-axis and short-axis images, with epicardial and endocardial contours used to quantify longitudinal, circumferential, and radial deformation[6]. Echocardiographic parameters for each trial are summarized in Tables S1–S3.

**Cardiac histology**

Hearts were immediately collected after sacrifice. Cardiac tips were snap-frozen at -80 °C for RNA isolation. The remaining tissue was either fixed in 10% neutral buffered formalin solution (Merck) for 36 h at room temperature (RT) and then switched to 1% until further use, or fixed in 4% paraformaldehyde (PFA) in PBS for 5 h at 4 °C.

For Picrosirius Red staining (PSR), formalin-fixed heart tissues were embedded in paraffin and transversally sectioned (2 µm) using an HM 340E Rotary Microtome (Epredia). Prior to staining with PSR, tissue sections were deparaffinated, rehydrated, washed and incubated in PSR working solution (Morphisto) according to manufacturer's protocol, with the exception of using ROTIClear (Carl Roth) as a xylol substitute. After washing and dehydration, samples were mounted in water-free medium (ROTIMount, Carl Roth, HP68.1).

For IHC stainings, PFA-fixed cardiac tissue was washed in PBS and incubated in 15% sucrose in PBS for 6-12h and 30% sucrose for 6-12h under constant rotation at 4°C for cryoprotection. Cardiac tissue was frozen embedded in Tissue-Tek O.C.T. Compound (Sakura Finetek) in -40°C methyl butane and stored at -80% until further use.  Hearts were cut into 7 µm cryosections on a Cryostat Microtome (CMC3050 S, Leica Biosystems). Prior to staining, cryosections were re-hydrated in Tris-buffered saline (TBS) for 10 min at RT and subsequently permeabilized in TBS with 0.1% Tween 20 (TBST) for 5 min at RT. Blocking was performed with 5% bovine serum albumin (BSA) and 10% normal donkey serum (NDS) in TBST for 20 min at RT.

For wheat germ agglutinin (WGA) staining, cryosections were incubated with WGA Fluorescein (1:800, Vector Laboratories) and DAPI staining solution (1:1000, Biotium) in TBST + 5% NDS for 2 h at RT. For extracellular matrix (ECM) protein staining, cryosections were incubated with primary antibodies, rabbit anti-fibronectin (1:100, polyclonal, abcam) and goat anti-type I collagen (1:50, polyclonal, Southern Biotech), in TBST + 5% NDS overnight at 4 °C. After washing in TBST, cryosections were incubated with secondary antibodies, donkey anti-rabbit IgG Alexa Fluor Plus 488 (1:1000, Thermo Fisher) and anti-goat IgG Alexa Fluor Plus 555 (1:1000, Thermo Fisher), together with DAPI (1:1000, Biotium) in TBST + 5% NDS for 1 h at RT. After washing in TBS, slides were mounted with DAKO fluorescent mounting medium.

Slides were imaged using the PANNORAMIC$^{TM}$ MIDI II digital slide scanner (3DHistech) with a 20x objective. For each heart, six representative snapshots were taken at 20x magnification using CaseViewer (v2.2.1, 3DHistech) for downstream analysis.

For PSR and ECM stainings, the percentage of positively stained area was quantified. Snapshots were pre-processed using Fiji to generate a binary image of stained versus unstained areas, and quantification was performed using CellProfiler. For WGA staining, the size of the unstained area was quantified. After pre-processing in Fiji, each snapshot was segmented in Ilastik by classifying pixels as stained or unstained using an interactive machine-learning algorithm. The size of segmented objects (unstained areas) was subsequently quantified using CellProfiler. The mean value across all snapshots was calculated per animal.

**Wire myography**

From a subset of animals (n=5 STNx, n=8 STNx + Abx), mesenteric arteries were removed, immediately transferred to cold (4°C) oxygenated (95% $O_2$/ 5% $CO_2$) physiological salt solution (PSS: 119 NaCl, 4.7 KCl, 1.2 $KH_2PO_4$, 25 $NaHCO_3$, 1.2 $Mg_2SO_4$, 11.1 glucose, and 1.6 $CaCl_2$ in mmol/L), and dissected into 2-mm rings (n=2 per animal). Perivascular fat and connective tissues were removed. Each ring was positioned between two stainless steel wires (0.0394-mm dia.) in a 5-mL PSS-filled organ bath of Small Vessel Myograph (DMT 630MA, Danish Myo Technology, Aarhus, Denmark). The PSS was continuously oxygenated and kept at 37°C (pH 7.4). Mesenteric rings were placed under a tension equivalent to that generated at 0.9 times the diameter of the vessel at 100 mm Hg. This normalization procedure was performed to obtain the passive diameter of the vessel at 100 mm Hg. The software LabChart 8 was used for data acquisition and display. The rings were precontracted with the KCl buffer (119 NaCl, 60 KCl, 1.2 $KH_2PO_4$, 25 $NaHCO_3$, 1.2 $Mg_2SO_4$, 11.1 glucose, and 1.6 $CaCl_2$ in mmol/L) and equilibrated until a stable resting tension was acquired. To test the endothelial function, vessels were precontracted with 1 µmol/L phenylephrine. After all vessel segments had reached a stable contraction plateau, increasing doses of acetylcholine were administered to the organ baths (3 to 1000 nmol/L). Tension is expressed as a percentage of the steady-state tension (100%) obtained with phenylephrine.

**Immunophenotyping**

<u>Plasma collection, immune cell isolation and flow cytometry measurements</u>

Whole blood was collected in lithium-heparin syringes (80 IU/3 mL, BD Vacutainer A-Line) for anticoagulation and centrifuged at 1,500 g for 15 min at RT. Plasma was collected and frozen at -80 °C, and the remaining blood cell pellet was resuspended in erythrocyte lysis buffer (83 g/L $NH_4Cl$, 8.5 g/L $NaHCO_3$, 10 mM EDTA) and incubated at 37 °C for 6 min for peripheral blood cell (PBC) isolation. After centrifugation at 400 g for 5 min (RT), supernatant was discarded and cells underwent a second erythrocyte lysis step (37 °C, 4-6 min). Reaction was stopped using ice-cold PBS + 0.5 % BSA + 2 mM EDTA (referred to as FACS buffer) and cell suspension was filtered through a 30 µm strainer.

Spleens were excised and kept in FACS buffer at 4 °C. Single cell suspensions were obtained by pressing tissue through 70 µm EasyStrainer, followed by erythrocyte lysis and filtration through 40 µm EasyStrainer.

Hearts were excised, atria were removed and tissue was kept in HBSS without $Ca^{2+}/Mg^{2+}$ + 10 mM HEPES (referred to HBSS w/o) at 4 °C. Ventricles were cut into 2-3 mm pieces before transfer into gentleMACS C Tubes (Miltenyi Biotec). A fresh digestion mix was prepared with DNase I (30 U/mL), Hyaluronidase I-S (60 U/mL), Collagenase XI (125 U/mL) in 1 mL HBSS with $Ca^{2+}/Mg^{2+}$. Samples were incubated for 1 h at 700 rpm and 37 °C, followed by mechanical dissociation using the gentleMACS Octo Dissociator (Miltenyi Biotec, program m_heart_02). Digestion was quenched using ice-cold PBS + 10 % FBS. Cells were pelleted (400 g, 10 min, 4 °C), treated with erythrocyte lysis buffer for 3 min at RT, neutralized with FACS buffer, and filtered sequentially through 70 µm and 40 µm EasyStrainers. An aliquot was collected for absolute counting (1:100 dilution).

Intestines were excised, flushed with HBSS w/o to remove all contents, divided into small intestine (duodenum, jejunum, ileum) and large intestine (cecum, colon) and stored in HBSS w/o at 4 °C. Lamina propria cells were isolated using the Lamina Propria Dissociation Kit, mouse (Miltenyi Biotec).

Heart and lamina propria cell suspensions were subjected to a 40%/80% Percoll density gradient (centrifugation at 800 g, 20 min, RT, no brake). Immune cells were harvested at the interphase and washed in FACS buffer.

Splenocyte and lamina propria cell suspensions were counted on a LUNA-FL Dual Fluorescence Cell Counter (Logos Biosystems), centrifuged and resuspended at $1x10^6$ cells per 100 µL medium (RPMI 1640 + 10% FBS + 1% penicillin/streptomycin [P/S, 10,000 U/mL / 10,000 µg/mL]) for further analysis. PBC and heart cell suspensions were directly centrifuged and resuspended in 100 µL medium per flow cytometry panel.

Cells were either immediately stained for flow cytometry or restimulated with 50 ng/mL PMA, 500 ng/mL ionomycin, and 0.75 µL/mL GolgiStop in medium for 4 h at 37 °C, 5 % $CO_2$. Viability was assessed with Live/Dead Fixable Aqua for 405 nm (Thermo Fisher), surface markers were stained in Brilliant Stain Buffer (BD Biosciences) with Fc block (Miltenyi Biotec), then cells were fixed/permeabilized (eBioscience FoxP3/Transcription Factor Buffer Kit) for intracellular staining. All antibody incubations were 30 min at 4 °C (Table S7). Data were acquired on a BD LSRFortessa with high throughput sampler (HTS) using BD FACSDiva and analyzed in FlowJo 10.8.1.

Flow cytometry analysis

Immune cell populations were identified by flow cytometry using the gating strategies in Figure S9. For the immune cell abundance analysis, batch effects arising from different flow cytometry acquisition days were corrected using ComBat from the *sva* R package (v.3.20). Raw abundance values were imported into R and the matrix of cell-type frequencies was extracted. Batch identifiers were supplied as the batch variable, and group assignment was included as a covariate in the ComBat model matrix. ComBat was run with parametric priors and without generating prior plots, yielding batch-adjusted abundance values that were re-combined with sample metadata for downstream analyzes. All subsequent analyzes were performed on the batch-corrected dataset using the same gating-based cell populations described above. UMAP visualizations were generated from the T-cell panel using two representative samples per group. $CD3^+$ T cells were isolated using the hierarchical gating strategy, and all gated events (10,000 $CD3^+$ T cells per sample) were used for dimensionality reduction. UMAP embeddings were computed in FlowJo using the UMAP_R plugin (version 4.1.1) with Euclidean distance, minimum distance = 0.1, and 15 nearest neighbours. FlowJo's default logical transformation was applied to all fluorescence channels. Group-level embedding intensity plots were created in FlowJo by overlaying all cells from the same group and visualizing them as smoothed density distributions.

**Mouse AhR reporter assay**

Mouse AhR-responsive reporter cell line (H1L1.1c2), derived from mouse hepatoma (Hepa1c1c7) cells and stably transfected with the pGudLuc1.1 vector containing AhR-responsive dioxin response elements (DRE) upstream of a luciferase gene, was kindly provided by Prof. Michael S. Denison, University of California, Davis, CA, USA. H1L1.1c2 cells were cultured in Opti-MEM (Thermo Fisher, 31985070) supplemented with 1% FBS (Merck, F7524), 1% P/S, and 0.2 mg/mL G418 (Sigma, A1720-1G), and maintained at 37 °C with 5% $CO_2$. Subculture was performed twice a week using TrypLE™ Express Enzyme (Gibco, 12604-02).

For the assay, cells were seeded in 96-well plates at a density of $7.5x10^4$ cells/well and allowed to attach for 24 h. Cells were then incubated with 10% mouse plasma in the same medium for 24 h. Assays were performed in triplicate, and 6-formylindolo[2-b]carbazole (FICZ) was included as a positive control at a concentration of 10 nM. After treatment, luciferase activity and cell viability were quantified using the One-Glo™ + Tox Luciferase Reporter and Cell Viability Assay (Promega, E7120) according to the manufacturer's instructions. Luminescence and fluorescence were measured using a microplate reader (Infinite 200 plate reader, Tecan). Luminescence values were normalized to cell viability (fluorescence signal) and subsequently expressed relative to the FICZ control within each plate.

**$T_H17$ polarization assay, murine**

Mesenteric and peripheral lymph nodes (LN) were harvested from 129S2/SvPasOrlRj mice and kept in cold FACS Flow buffer. LN were mechanically dissociated through a 70 µm cell strainer using cold FACS Flow buffer. The cell suspension was subsequently filtered through a 40 µm strainer, washed with 20 mL FACS Flow buffer, and centrifuged at 400 × g for 10 minutes at 4°C. Cells were resuspended in FACS Flow buffer at 1 × 10^7 cells per 100 µL.

CD4+ T cells were enriched via magnetic-activated cell sorting (MACS) using the MojoSort Mouse CD4 T Cell Isolation Kit (BioLegend) with minor modifications for the use of magnetic columns. Briefly, cells were incubated with a biotinylated antibody cocktail (2 µL per 10^7 cells in 100 µL) for 15 minutes at 4°C, followed by Streptavidin Nanobeads (2 µL per 10^7 cells) for an additional 15 minutes at 4°C. Cells were separated using LS columns (Miltenyi Biotec) and the unlabelled fraction was used for further analysis. For naïve CD4+ T cell isolation, cells were stained with CD4 (VioGreen), CD25 (PE), CD44 (BB700), CD62L (BV421), and a live/dead stain (Zombie Green) and $CD4^+CD25^-CD44^{low}CD62L^{high}$ cells were sorted on a FACS Melody (BD Bioscience, 100 µm nozzle).

96-well flat-bottom plates were coated overnight at 4°C with anti-CD3 antibody (2 µg/mL in PBS, 50 µL per well). Naïve CD4+ T cells were resuspended in $T_H17$ differentiation medium (RPMI-1640 supplemented with 10% FCS, 1% penicillin-streptomycin, 1% L-glutamine, 1% non-essential amino acids, 1 mM sodium pyruvate, and 0.01 mM β-mercaptoethanol). Cells were plated at 5 × 10^4 cells per well in a final volume of 200 µL with anti-CD28 (2 µg/mL), mIL-6 (80 000 U/mL, Miltenyi Biotec), rhTGFβ1 (40 U/mL, Miltenyi Biotec) and mIL-1β (33 600 U/mL, Miltenyi Biotec) and test compounds (IxS, KCl, AhRi at concentrations indicated in the respective figure) or DMSO vehicle control, as appropriate. Cells were cultured at 37°C with 5% $CO_2$ for 4 to 5 days and subsequently restimulated for 4 hours with PMA (50 ng/mL), ionomycin (750 ng/mL), and GolgiPlug (0.5 µL/mL). Following restimulation, cells were washed with FACS Flow buffer, stained with BD Horizon Fixable Viability Stain 700 (1:500 dilution) for 30 minutes cold, fixed, and permeabilized using Fix/Perm buffer (eBioscience), and stained with the antibodies indicated in Table S7. Cells were analyzed using a BD LSRFortessa with high throughput sampler (HTS). Data were processed using FlowJo.

**RNA isolation**

Splenocytes were isolated as described above, enriched for CD45+ using mouse CD45 MicroBeads (Miltenyi Biotec). Hearts were excised and apex was snap frozen at -80°C. Spleens were excised and a small piece of tissue was placed into 300 µL RNAprotect Tissue Reagent (Qiagen) at 4 °C for a minimum of 24 h. RNAprotect Tissue Reagent was then discarded, and tissues were frozen at -80 °C.

RNA isolation from splenocytes and spleen was performed using the RNeasy Mini Kit (Qiagen). RNA isolation from heart was performed using the RNeasy Fibrous Tissue Mini Kit (Qiagen). For cell homogenisation, enriched CD45+ splenocytes were resuspended in buffer RLT + β-mercaptoethanol and vortexed vigorously. Tissue pieces were immersed in RLT buffer + β-mercaptoethanol with 1.5 mm ceramic beads and placed into a Precellys 24 tissue homogeniser. RNA was subsequently isolated from both sample types according to the instructions of the respective kits.

Fibroblasts on well plates were washed in PBS and incubated with 700 µL Qiazol Lysis Reagent (Qiagen) per well at -80 °C over night. Then, lysed cells were harvested, transferred to a tube and vortexed vigorously. 200 µL of chloroform was added and samples were centrifuged at 12,000 g for 15 min at 4 °C. Afterwards, the aqueous phase was collected and RNA was isolated with the RNeasy Mini Kit according to the manufacturer's protocol.

**Quantitative polymerase chain reaction (qPCR)**

All qPCR was performed on a QuantStudio™ 5 Real-Time PCR System (Thermo Fisher Scientific) using a 384-well plate format. For each 5-µL reaction mix, 4 ng of DNA or cDNA was used, and all detections were performed in triplicate. All primer and probe sequences and concentrations are listed in Table S6.

For all SYBR-Green based detection, the PowerUP™ SYBR™ Green Master Mix (Thermo Fisher Scientific) was used with a fast cycling protocol of 50 °C for 2 min, 95 °C for 2 min, followed by 40 cycles of 95 °C for 1 s and 60 °C for 30 s. For all TaqMan-based detection, the TaqMan® Multiplex Master Mix (Thermo Fisher Scientific, 4461882) was used with a fast cycling protocol of 95 °C for 20 s, followed by 40 cycles of 95 °C for 2 s and 60 °C for 20 s.

For *16S* quantification, cecal DNA was amplified and detected using SYBR Green and the standard curve method. Standard curves were generated using 10-fold serial dilutions ranging from $10^1$ to $10^9$ copies of the *16S* rRNA gene of *E. coli* (Invitrogen) amplified with primers 27F (5′-GTTTGATCCTGGCTCAG-3′) and 1492R (5′-CGGCTACCTTGTTACGAC-3′). Using the resulting curve, *16S* rDNA copy numbers per gram content were calculated.

For all other genes, cDNA was synthesized prior to qPCR using the High Capacity cDNA Reverse Transcription Kit (Thermo Fisher Scientific). Murine *Col1a2*, *Acta2*, *Cyp1a1* and *18s* were detected using TaqMan. Gene expression was normalized to the *18s* signal using the ΔΔCt method. All other murine genes were quantified using SYBR Green, and *Hprt* or *Gapdh* as the housekeeping genes. All human genes were quantified using SYBR Green and normalized to *HPRT1* using the ΔΔCt method. All primer and probes are shown in Table S6.

**Bulk mRNA sequencing**

Illumina stranded mRNA library preparation and bulk mRNA sequencing were performed at the BIH/MDC Genomics Technology Platform. Paired-end sequencing was performed on an Illumina NovaSeq X Plus platform, targeting ~30 million reads per sample. Transcript quantification of raw sequencing data to the *Mus musculus* reference transcriptome (GRCm38) was performed using Salmon (v1.10.1). Quality assessment of the preprocessing was performed using MultiQC (v1.6). Following quality control, 26.4M-34.7M paired-end reads per sample were retained and included in downstream analysis. Gene symbols were generated from Ensembl gene IDs using biomaRt (v2.52.0, Ensembl release 102). Post-processing quality control and downstream data analysis were conducted using DESeq2 (v1.42.1).

For the Abx trial, separate DeSeq2 datasets were generated for each reference condition (Sham, STNx). Lowly expressed genes were removed by filtering for transcripts with at least 1 count per million in a minimum of two samples using edgeR (v4.0.16). Differential expression analysis was performed using the DeSeq2 pipeline. $Log_2$ fold changes were adjusted using lfcShrink with the ashr method (v2.2.63), and p-values were corrected using a false discovery threshold of 10%. Variance-stabilized expression values were obtained using the rlog transformation.

Gene set enrichment analysis (GSEA) was performed on the overlap of significantly regulated genes from the STNx vs. Sham and STNx Abx vs. STNx comparisons (adjusted p-value < 0.1). Curated *Mus musculus* gene sets were downloaded from MSigDB. Enrichment was calculated with Fisher's exact test and FDR correction, using all Ensembl gene IDs present in the complete MSigDB *Mus musculus* dataset as the background. Pathways and GO terms not relevant to the study were excluded from visualisation using targeted keyword filtering.

For the AhRi trial, differential expression analysis with DeSeq2 was restricted to the set of DEGs in STNx that were Abx responsive. Batch effects between experimental rounds were corrected using ComBat-seq (sva, v3.50.0). Genes significantly regulated by AhRi-treatment in the same direction as the Abx-dependent response (p < 0.1) were subjected to GSEA.

For both trials, targeted gene set variation analysis (GSVA) was performed on rlog-transformed expression values using curated gene sets relevant to the $T_H17$ axis (gsva package, v1.50.5).

**Single-cell RNA-sequencing**

We analyzed publicly available mouse cardiac single-nucleus and human single-cell RNA-seq datasets[7,8]. Processed Seurat objects were imported into R (v4.2.2) using Seurat (v5.0). No reclustering or reannotation was performed. For each dataset, cells belonging to diseased samples (mouse CKD or human CKD) and appropriate controls were subset for downstream analyzes. Resident structural cell types were selected based on the original annotations, and expression of IL17ra was quantified from log-normalized RNA values to identify positive and negative stromal populations. Extracellular matrix (ECM) activity was assessed using an extended NABA matrisome gene list[9] and compared using Wilcoxon tests. AhR pathway activity was similarly scored using a recently published AhR-responsive gene set[10]. Differential expression was performed using the MAST framework[11], including only genes expressed in more than 10% of nuclei. Genes were considered differentially expressed when meeting: adjusted $p < 0.1$, absolute $\log_2$ fold change > 0.3, and average normalized expression > 3 in Il17ra$^+$ cells. Differentially expressed genes were compared with curated fibroblast transcriptional signatures derived from human cardiac and stromal datasets[12] to identify enriched transcriptional programs. Briefly, DEG were intersected with each fibroblast signature and the number of overlapping genes, proportion of signature genes present among upregulated genes, and average FC and expression of the overlapping genes were used to find similar transcriptional programs. Cytokine-response enrichment was conducted from DEG of cardiac lymphocytes as described above using the Immune Response Enrichment Analysis (IREA)[13] software.

**Murine cardiac fibroblasts**

Hearts were freshly harvested from 7-10 weeks old male and female C57BL/6J mice and rinsed twice in PBS. Aortic and atrial tissues were removed, and ventricles were minced and transferred into 1 mL of enzymatic digestion solution containing DNase I (300 U/mL), Hyaluronidase I-S (60 U/mL), Collagenase XI (125 U/mL), and Collagenase I (450 U/mL) in DMEM. Digestion was performed on a shaking incubator (800 rpm, 37 °C) for 45 minutes.

Following digestion, the cell suspension was filtered through a 70 µm strainer using PBS and centrifuged at 400 g for 10 minutes at 37 °C. The pellet was resuspended in 1 mL erythrocyte lysis buffer (Erylysis), incubated for 3 minutes at RT, and quenched with PBS. Cells were centrifuged again under the same conditions, supernatant was discarded, and the pellet was resuspended in 4 mL fibroblast culture medium (DMEM supplemented with 20% FBS, 2% penicillin-streptomycin, 1 ng/mL endothelial cell growth factor [ECGF], and 10 µg/mL heparin). Cells were maintained at 37 °C and 5% $CO_2$. Media was changed after 2.5 h and further replaced every 2-3 days. For subculture, fibroblasts were detached using Trypsin-EDTA (1mL, 2-3 min incubation). Reaction was stopped with culture media, fibroblasts were transferred to a 15 mL tube and centrifuged at 400 g for 10 min. Pellet was resuspended in culture media, counted and plated into tissue flasks or well plates depending on cell counts (6.000-10.000 cells/cm$^2$).

Assay was performed on a 6-well plate. In brief, fibroblasts were starved in culture media containing 1% FBS and incubated for 4 h at 37 °C, followed by a 24 h incubation with the test substances. Murine recombinant IL-17A was used at 1 ng/mL in culture media. Potassium 3-indoxyl sulfate was used at 50 µM and 100 µM on its own, and at 100 µM in combination with IL-17A in culture media. After 24 hours incubation, fibroblasts were washed in PBS and incubated with 700 µL Qiazol Lysis Reagent per well at -

80 °C over night. Lysed fibroblasts were then harvested, RNA was isolated and RT-qPCR was performed as described above.

For immunofluorescence staining, 18x18 mm coverslips were sterilized with ethanol 70% and placed into 6-well plates. Fibroblasts were grown on the coverslips in culture media until desired confluence. Cells were then washed with PBS and fixed with 4% Paraformaldehyde, followed by permeabilization with 0.1% Triton X-100 and blocking with 10% normal donkey serum (NDS) in PBS.

For PDGFR$\alpha$ and VIM staining, cells were incubated with conjugated antibodies (anti-mouse CD140a PE, 1:100, clone APA5, Miltenyi, 130-102-502 and AF647 anti-vimentin, 1:500, clone EPR3776, ab194719) for 2 h at RT in PBS + 10% NDS. For COL-1 and IL17RA staining, cells were incubated with primary antibodies (goat anti-type I collagen, 1:500, polyclonal, Southern Biotech, 1310-01 and rabbit anti-IL17RA, 1:500, polyclonal, abcam, ab180904) for 2 h at RT in PBS + 10% NDS. After washing with PBS + 1% NDS, secondary antibodies, donkey anti-rabbit IgG Alexa Fluor Plus 488 (1:1000, Thermo Fisher) and anti-goat IgG Alexa Plus 555 (1:1000, Thermo Fisher), were incubated together with DAPI (1:1000, Biotium) in PBS + 10% NDS for 1 h at RT. After washing, coverslips were taken out of the well plates and mounted onto microscopy slides using DAKO fluorescent mounting medium. Slides were imaged on the Keyence Compact Fluorescence microscope with the Plan Apochromat 20x objective lens.

For flow cytometry staining, cells were stained for PDGFR$\alpha$, CD45 and CD31 (antibody concentrations in Table S7) together with SYTOX™ Green Dead Cell Stain (1:500, Invitrogen, S34860) and Fc blocking reagent (Miltenyi Biotec, 130-092-575) in FACS buffer. Data were acquired on a BD LSRFortessa using BD FACSDiva and analyzed in FlowJo 10.8.1.

**Splenocyte incubation assay**

Spleens from 129S2/SvPasOrlRj mice were processed to obtain spleen single-cell suspensions as described above. For *ex vivo* stimulation, splenocytes were incubated for 4 h at 37 °C, 5 % $CO_2$ in RPMI-1640 in the presence of IxS and AhRi at the indicated concentrations. After incubation, cells were resuspended in buffer RLT + $\beta$-mercaptoethanol and frozen at −80 °C for subsequent RNA isolation as described above.

**Human luciferase reporter assay**

Human AhR reporter cell line (HT29-Lucia™ AhR, InvivoGen, ht21-ahr), containing AhR-responsive DRE upstream of a Lucia luciferase gene, was cultured in Dulbecco's Modified Eagle Medium (DMEM) supplemented with 10% FBS and 100 µg/mL Zeocin (InvivoGen, ant-zn-05), maintained at 37 °C with 5% $CO_2$. Cells were subcultured once per week using a standard Trypsin-EDTA protocol and partially replenished with fresh medium mid-week between passages.

For the assay, 180 µL of cell suspension per well was seeded on a 96-well plate at a density of $2.8 \times 10^5$ cells/mL in medium without Zeocin. For the dose-response curve of AhRi, cells were incubated with 100 µM IxS and AhRi concentrations between 1 nM and 10 µM or VEH (vehicle, DMSO content of the highest AhRi concentration) for 24 h in quadruples. To assess the AhR activation potential of human biomaterial, cells were incubated with 10% patient serum and AhRi in a final concentration of 1 or 0.1 µM, or VEH for 24 h in triplicates. For quantification, 20 µL of culture supernatant was used to determine luciferase activity using the QUANTI-Luc™ Luciferase Detection Reagent (InvivoGen, rep-qlc4lg1). Luminescence was measured using a microplate reader (Infinite 200 plate reader, Tecan).

**Human $T_H17$ polarization assay**

Peripheral blood mononuclear cells (PBMC) were collected together with serum from healthy volunteers under the ethics approval described above (EA2/162/17). PBMC were isolated from EDTA-anticoagulated blood of a healthy donor by diluting samples 1:1 in PBS + 2% FBS and layering them into 50 mL SepMate tubes (Stemcell) prefilled with 15 mL Pancoll (PAN Biotech). After centrifugation at 1200 g for 10 min at RT, the mononuclear cell layer was collected, transferred into a fresh 50 mL tube, and washed with PBS + 2% FBS, followed by erythrocyte lysis.

T cells were isolated from PBMC using MojoSort Human CD4 T Cell Isolation Kit (Biolegend) with LS columns (Miltenyi) following the manufacturer's protocol. To obtain naïve CD4 cells, the enriched T helper cell fraction was sorted using a BD FACSAria III (BD Bioscience, 70 µm nozzle) by staining for CD4, CD25, CD127, CD45RO (Table S8), and dead cells exclusion with DRAQ7 (1:250, Miltenyi Biotec).

Naïve CD4 T cells were resuspended in X-VIVO 15 serum-free hematopoietic media (Lonza Bioscience) and plated at $5 \times 10^4$ cells per well in anti-CD3-coated (10 µg/mL, BD) 96-well U-bottom plates and 1 µg/mL anti-CD28 (BD). To induce $T_H17$ polarization, naïve CD4 T cells were cultured with 12.5 ng/mL IL-1β, 25 ng/mL IL-6, 25 ng/mL IL-21, 25 ng/mL IL-23, and 5 ng/mL TGF-β (all Miltenyi Biotec), while $T_H0$ control wells were left without cytokine supplementation. Cells were treated with 2.5% serum from either hemodialysis patients or healthy controls in triplicates and maintained for 7 days at 37°C and 5% CO2.

Prior to analysis, cells were restimulated for 4 h with 10 ng/mL PMA (Merck), 500 ng/mL ionomycin (Merck), and 1 µL/mL GolgiPlug (BD), stained with LIVE/DEAD fixable aqua dye (Invitrogen), fixed and permeabilized using the Foxp3/Transcription Factor Staining Buffer Set (Invitrogen), and finally stained with antibody panels with Fc Block, CD4 and IL-17A (antibodies listed in Table S8).

Cells were analyzed using a BD LSRFortessa with HTS and data were processed using FlowJo (v10.10).

**Human cardiac fibroblasts**

Human cardiac fibroblasts were obtained from PromoCell (Germany). Cells were seeded (4000 cells per $cm^2$ seeding density) with 18 mL of Fibroblast Growth Medium 3 (PromoCell, Germany) into a tissue flask T75. Media contained supplements (PromoCell, Germany), P/S (1%) and Amphotericin B (Gibco, 0.2%). Cells were maintained at 37 °C and 5% $CO_2$ with medium exchanged every 2-3 days and passaged at 70-90% confluence as described above. Subculture was performed using the standard Trypsin-EDTA protocol.

For all experiments, cells were seeded at manufacturer recommended densities and allowed to attach overnight. The following day, cultures were starved for 4h in medium containing 1% FBS and subsequently stimulated for 24h under the appropriate conditions. Treatments included IL-17A (1 ng/mL), IxS (100 µM), AhRi (1 µM), Secukinumab (Cosentyx, 10 nM), human serum (HD/HC; 10%), or the corresponding combinations. After stimulation, cells were washed once with PBS and lysed in QIAzol as descrived above. Lysed cells were then harvested, RNA was isolated and RT-qPCR was performed as described above.

To assess viability and proliferation, fibroblasts were co-incubated with IL-17A and IxS in starved medium for 24 h. After detaching, cells were stained with Zombie Green Fixable Viability Kit (1:500, BioLegend), fixed/permeabilized (eBioscience FoxP3/Transcription Factor Buffer Kit) for intracellular staining, and stained for Ki-67 (Table S8.)
